# Genome-wide evidence for cryptic multi-generational introgression between invasive *Bursaphelenchus xylophilus* and native *B. mucronatus*

**DOI:** 10.64898/2026.09.18.752818

**Authors:** Yuzuki Ikeda, Naoko Ishikawa, Kenichi Yanagisawa, Yoshihisa Suyama, Ryoji Shinya

## Abstract

Hybridization between invasive and native species can generate novel genetic variation with ecological and evolutionary consequences, yet introgression often remains cryptic. We investigated natural hybridization between the invasive pinewood nematode *Bursaphelenchus xylophilus*, the causal agent of pine wilt disease, and native *Bursaphelenchus mucronatus* in Japan. Using multiplexed inter-simple sequence repeat genotyping by sequencing, we genotyped 342 field-collected nematodes and then conducted pedigree-calibrated analysis using HIest v2.0, based on 48 highly informative diagnostic single-nucleotide polymorphisms (SNPs), followed by ADMIXTURE, complementary NeighborNet and principal component analyses of a broader 2,756-SNP dataset. The results identified nine high-confidence hybrids (2.6%) across two consecutive years (2023–2024), including relatively balanced F1-like individuals and strongly parentally biased advanced backcross-like individuals, demonstrating recurrent, multigenerational introgression. Four hybrids (44.4%) were undetectable using a single-locus diagnostic marker, showing that advanced introgression can escape conventional surveillance. Chemotaxis assays detected no volatile-mediated pre-mating discrimination, whereas post-mating barriers were incomplete and asymmetric. Crosses with *B. xylophilus* as the maternal parent showed reduced hatchability, particularly with *B. mucronatus mucronatus*, while reciprocal crosses generally retained higher hatchability, although some showed reduced fecundity. These results provide a mechanistic basis for natural gene flow in a recent secondary contact zone. Because *B. mucronatus kolymensis* is widespread in Europe and produces viable hybrid offspring with *B. xylophilus* in both directions, pathogen establishment could create new opportunities for introgression. Our findings reveal cryptic, multigenerational introgression between an invasive forest pathogen and its native congener, highlighting the importance of genomic surveillance for forest disease management and invasion-risk assessment under climate change.

## Introduction

Interspecific hybridization between invasive and native species is increasingly recognized as a critical driver of rapid evolutionary change and ecological disruption. When closely related species come into secondary contact following biological invasion, gene flow through hybridization can create novel genetic combinations that may enhance adaptive potential beyond that of either parental species (Abbott et al., 2013; Ellstrand & Schierenbeck, 2000). Such hybridization events have been documented across diverse taxa, from plants (e.g., *Spartina* cordgrasses; Ainouche et al., 2009) to insects (e.g., *Heliconius* butterflies; Nadeau et al., 2012) and mammals (e.g., *Mus* mice; Janoušek et al., 2012). Critically, hybrids can exhibit transgressive traits—phenotypes that exceed the range of both parents—potentially facilitating range expansion into environments where neither parent thrives (Rieseberg et al., 1999). Furthermore, repeated backcrossing between hybrids and parental species leads to introgressive gene flow, which can alter the genetic architecture of recipient populations and enable colonization of novel ecological niches (Arnold & Martin, 2010). Therefore, understanding the frequency, direction, and evolutionary consequences of natural hybridization is essential for predicting and managing the ecological impacts of biological invasion.

Interaction between the invasive pinewood nematode *Bursaphelenchus xylophilus* (Steiner & Buhrer, 1934) Nickle and the native *Bursaphelenchus mucronatus* Mamiya & Enda provides a useful system for investigating natural hybridization dynamics. *Bursaphelenchus xylophilus*, the causal agent of pine wilt disease, was likely introduced to Japan from North America in the early 1900s through infected timber (Jikumaru & Togashi, 2008; Zhao et al., 2008). Following its establishment, the disease front has expanded progressively northward across the Japanese Archipelago, causing extensive mortality in susceptible pine species (Zhao et al., 2008). This nematode has subsequently spread to other regions, including China, South Korea, and parts of Europe, representing a major threat to global pine forest ecosystems (Arbuzova et al., 2025; Zhao et al., 2008). *Bursaphelenchus mucronatus*, which is native to Eurasia including Japan, shares a similar life history, as both species are vectored by *Monochamus* longhorn beetles and colonize pine xylem tissues, but it exhibits negligible or weak pathogenicity to pine hosts (Mamiya & Enda, 1979). In Japan, *B. xylophilus* is mainly dispersed by *Monochamus alternatus* (Mamiya & Enda, 1972; Morimoto & Iwasaki, 1972), whereas *B. mucronatus* is primarily dispersed by *Monochamus saltuarius*, which exhibits segregated distribution in high-altitude pine forests (Ozawa et al., 2021). Importantly, climate warming is enabling *B. xylophilus* to advance into cooler northern regions (Xiao et al., 2024), increasing its geographic overlap with *B. mucronatus* populations and creating conditions conducive to interspecific hybridization. Recent evidence from China has confirmed the occurrence of natural hybrids between these species based on species-specific polymerase chain reaction (PCR) primers (Li et al., 2021), raising critical questions about the extent and consequences of hybridization in other parts of the invaded range.

Although laboratory studies have demonstrated that *B. xylophilus* and *B. mucronatus* can produce viable hybrids under controlled conditions (Taga et al., 2011), and that some hybrid lineages exhibit enhanced pathogenicity or altered host ranges (Bolla & Boschert, 1993; Togashi et al., 2023), our understanding of natural hybridization remains limited by methodological constraints. Previous detection efforts have relied exclusively on single-locus diagnostic markers (Li et al., 2021; Matsunaga et al., 2019), which are useful for identifying F1 hybrids. However, when marker loci become homozygous through segregation in F2 hybrids or successive backcrossing in later generations, advanced-generation hybrids become indistinguishable from pure parental species. Such cryptic hybridization can lead to severe underestimation of gene flow, obscuring the true extent of introgression in natural populations (Anderson & Thompson, 2002; Vähä & Primmer, 2006). Genome-wide approaches overcome this limitation by providing sufficient statistical power to detect even the subtle admixture signatures characteristic of advanced backcrosses (Gompert & Buerkle, 2016). Among these, multiplexed inter-simple sequence repeat (ISSR) genotyping by sequencing (MIG-seq) involves the generation of genome-wide single-nucleotide polymorphism (SNP) data via PCR-based next-generation sequencing, without requiring a high-quality reference genome (Suyama & Matsuki, 2015). Although MIG-seq has proven effective for detecting hybridization between closely related plant species (Suetsugu et al., 2021; Tamaki et al., 2016; Watanabe et al., 2018), to our knowledge, it has not yet been applied to detect backcrossed hybrid generations in wild animal populations.

In this study, we conducted MIG-seq-based SNP analyses to determine whether hybridization between invasive *B. xylophilus* and native *B. mucronatus* is ongoing in natural populations in Japan, whether introgression extends beyond the F1 generation, and which components of reproductive isolation may constrain interspecific gene flow. We first established an empirically calibrated HIest framework using laboratory-generated parental, F1, B1, and B2 individuals of known pedigree. Then, we genotyped 342 nematodes collected from *M. alternatus* vectors in Shiojiri, Nagano Prefecture, a recent secondary contact zone where both species co-occur, over two consecutive years (2023–2024). Hybrid assignments based on diagnostic SNPs were further evaluated using ADMIXTURE software and complementary analyses of a broader genome-wide SNP dataset. In parallel, we quantified volatile-mediated sexual attraction, egg production and hatchability between *B. xylophilus* and two subspecies of *B. mucronatus*: *B. mucronatus mucronatus*, which occurs in East Asia including Japan, and *B. mucronatus kolymensis*, which is widespread in Europe (Braasch et al., 2011). By integrating genomic evidence from natural populations with experimental analyses of reproductive isolation, we aimed to clarify the extent and mechanisms of ongoing introgression in this invasive–native species complex and its implications for pine wilt disease management.

## Materials and methods

### Nematode strains and production of experimental hybrids

We used 10 strains of *B. xylophilus*, three strains of *B. m. mucronatus*, and six strains of *B. m. kolymensis* as parental reference strains (Table S1). All strains were maintained at 25°C on malt extract agar inoculated with *Botrytis cinerea*.

A subset of these strains was used to generate experimental hybrids of known pedigree for the calibration of hybrid ancestry analyses: four *B. xylophilus* strains (T-4, TBx#1, Ka4, and S10), three *B. m. mucronatus* strains (Un-1, NG-1, and Ioujima), and two *B. m. kolymensis* strains (TCS00 and Srf). Virgin females and males were isolated and crossed following Shinya et al. (2014). F1 progenies were individually reared to adulthood, and first- (B1) and second- (B2) generation backcrosses were generated by sequential backcrossing to the parental strains. These pedigree-confirmed hybrids were genotyped by MIG-seq and used as reference classes for hybrid ancestry estimation (Table S2).

### Field sampling, nematode extraction and species screening

Field sampling was conducted at the Nagano Prefectural Forestry Research Center in Shiojiri, Nagano Prefecture, Japan (36.143294°N, 137.995228°E; 850 m a.s.l.). Pine wilt disease was first recorded in Nagano Prefecture in 1981 and reached Shiojiri by 2014 (Yanagisawa et al., 2023). Notably, a previous survey in Nagano Prefecture recovered both *B. xylophilus* and *B. mucronatus* from the same *M. saltuarius* individual (Ozawa et al., 2021), demonstrating that both nematode species can co-occur within a single vector in this region, providing opportunities for interspecific contact.

Dead *Pinus thunbergii* trees were cut into logs (length, ∼100 cm; diameter, 5–30 cm) and maintained in outdoor emergence cages at the Nagano Prefectural Forestry Research Center and Meiji University. Emerging *M. alternatus* adults were collected daily from June 10 to August 12, 2023 (n = 115) and from June 10 to August 6, 2024 (n = 126).

Nematodes were extracted separately from individual beetles using a modified Baermann funnel method (Maehara et al., 2020). Each beetle-derived nematode assemblage was treated as a single population for subsequent analyses. For initial species screening, approximately 100 nematodes were randomly selected from each population, pooled for DNA extraction, and screened using diagnostic PCR markers for *B. xylophilus* and *B. mucronatus* (Li et al., 2021; Matsunaga et al., 2019; Matsunaga & Togashi, 2004; Table S3). Populations in which both species were detected were selected for individual-level genotyping. Detailed DNA extraction and PCR conditions are provided in the Supplementary Methods.

From each mixed-species population, 96 individual nematodes were randomly isolated into separate PCR tubes and subjected to DNA extraction. Based on DNA quality and quantity, 10–21 individuals per beetle-derived population were selected for MIG-seq analysis, yielding 170 and 172 field-collected samples in 2023 and 2024, respectively. Laboratory parental strains and laboratory-generated F1, B1, and B2 hybrids of known pedigree were included as reference samples for ancestry analyses.

As reference controls, 1–3 individuals from each of the 19 laboratory parental strains (10 *B. xylophilus*, three *B. m. mucronatus*, and six *B. m. kolymensis* strains; Table S1), together with laboratory-generated F1, B1, and B2 hybrids of known pedigree, were included in the MIG-seq analysis.

### MIG-seq genotyping and SNP discovery

Genome-wide SNP data were generated by MIG-seq (Suyama et al., 2022; Suyama & Matsuki, 2015). MIG-seq libraries were prepared separately for three sample sets: laboratory-generated hybrids of known pedigree and their parental individuals (n = 234), field-collected samples from 2023 (n = 170), and field-collected samples from 2024 (n = 172). Each sample set was processed independently to minimize potential batch effects among laboratory and field samples.

Library concentrations were initially assessed using a Qubit fluorometer and subsequently quantified by quantitative PCR (qPCR). All three MIG-seq libraries were sequenced using 80-bp paired-end reads on an Illumina MiSeq platform (Illumina, San Diego, CA, USA). All three MIG-seq libraries were also sequenced with 100-bp paired-end reads on a DNBSEQ-G400 platform (MGI, Shenzhen, Guangdong, China). Detailed MIG-seq PCR conditions, primer sequences, adapter sequences, and library preparation procedures are provided in the Supplementary Methods.

Raw reads were quality filtered using Trimmomatic v0.39 (Bolger et al., 2014), with platform-specific trimming parameters to account for read length differences. Both paired and unpaired reads that passed quality filtering were retained. Quality-filtered reads from the MiSeq and DNBSEQ-G400 platforms were then combined for each individual prior to SNP discovery and downstream analyses. Samples represented by fewer than 50,000 combined reads after quality filtering were excluded from subsequent analyses. All trimming parameters are provided in the Supplementary Methods.

The combined quality-filtered reads were assembled using ipyrad v0.9.102, against *B. xylophilus* reference genome (GCA_904066235.2_BXYJv5). Separate SNP datasets were generated for *B. xylophilus*–*B. m. kolymensis* and *B. xylophilus*–*B. m. mucronatus* comparisons. The maximum heterozygosity per locus was set to 0.6, and loci were required to be present in at least 76 of 108 individuals in the *B. xylophilus*–*B. m. kolymensis* dataset and at least 305 of 436 individuals in the *B. xylophilus*–*B. m. mucronatus* dataset. These thresholds were selected to balance locus retention and missing data across both datasets.

### HIest-based hybrid ancestry estimation and classification

Diagnostic SNP panels were constructed using balanced parental reference sets. For the *B. xylophilus*–*B. m. kolymensis* comparison, 16 individuals from each parental taxon were selected (n = 32), whereas 14 individuals from each parental taxon were used for the *B. xylophilus*–*B. m. mucronatus* comparison (n = 28). Because *B. xylophilus* was represented by multiple laboratory strains, individuals were subsampled to maintain balanced parental representation while capturing genetic variation among the major *B. xylophilus* lineages.

Diagnostic SNPs were identified from parental samples using PLINK v2.00a and v1.9. SNPs were retained if they met the following criteria: ≤ 25% missing data, with comparable call rates between parental taxa; allele-frequency divergence between parental taxa of |Δ| ≥ 0.80; at least one minor allele; and a minimum physical separation of 1 kb to reduce linkage among loci. These criteria initially yielded 43 and 58 diagnostic SNPs for the *B. xylophilus*–*B. m. kolymensis* and *B. xylophilus*–*B. m. mucronatus* datasets, respectively.

An additional filtering step was subsequently applied to the complete genotype matrices containing parental, laboratory-generated hybrid, and, for the *B. xylophilus*–*B. m. mucronatus* dataset, field-collected individuals. SNPs with > 20% missing data were removed, and loci showing identical genotype patterns across individuals were reduced to a single representative SNP. The final datasets contained 30 diagnostic SNPs for the *B. xylophilus*–*B. m. kolymensis* comparison and 48 diagnostic SNPs for the *B. xylophilus*–*B. m. mucronatus* comparison. Alleles were polarized such that the frequency of the *B. xylophilus* allele was greater than or equal to that of the alternative parental taxon.

Hybrid ancestry was estimated using HIest v2.0. For each individual, HIest estimated a hybrid index value (S), representing the proportion of *B. xylophilus* ancestry, and an interspecific heterozygosity value (H), representing the proportion of heterozygous diagnostic loci. Missing genotypes were excluded on a per-individual basis. On average, approximately 27 diagnostic SNPs (range 18–30) and 41 SNPs (range 2–48) were available per individual in the *B. xylophilus*–*B. m. kolymensis* and *B. xylophilus*–*B. m. mucronatus* datasets, respectively. Individuals represented by fewer than 10 diagnostic SNPs were excluded from class interpretation. The final HIest analyses included 108 and 429 individuals, respectively, parameter estimation was performed using simulated annealing optimization.

Laboratory individuals of known pedigree, including parental, F1, B1, and B2 classes, were used to establish empirical reference distributions in S–H space. For each pedigree class, empirical centers and standard deviations (SDs) of S and H were calculated. The classification performance of the theoretical and empirical approaches was evaluated using laboratory individuals of known pedigree that were independent of the parental reference individuals used for HIest estimation. Predicted classes were compared with known pedigree classes. For the empirical approach, classification accuracy was evaluated by leave-one-out cross-validation (LOOCV), in which each validation individual was classified using empirical class centers and SDs calculated after excluding that individual. Field-collected individuals were assigned to the closest reference class using class-specific standardized Euclidean distances in S–H space. For each reference class, deviations of an individual’s S and H values from the corresponding empirical class means were divided by the class-specific SDs of S and H, respectively. The standardized deviations were then combined as the Euclidean distance, and the class with the smallest distance was taken as the primary empirical assignment. The second-closest class and the difference between the second-smallest and smallest distances (distance margin) were also recorded to indicate the degree of separation between alternative assignments. Individuals of known pedigree retained their predefined class labels and were not reassigned. Only reference classes represented by pedigree-confirmed laboratory individuals were formally included in the assignment procedure.

Because validation indicated overlap among some parental and backcross classes, an additional sensitivity analysis was performed using ordinary Euclidean (OE) distance. OE distances were calculated from the same empirical class centers but without scaling the deviations in S and H by their class-specific SDs. Thus, the primary empirical classification accounted for class-specific variation in S and H, whereas the OE sensitivity analysis reflected only geometric proximity to the empirical class centers. OE assignments were used as a sensitivity check for field-sample interpretation rather than as an independent primary classification method.

For comparison, theoretical S–H positions expected under Mendelian inheritance were also calculated for each hybrid class. These theoretical expectations were used to visualize and evaluate deviations from idealized hybrid classes, whereas field individuals were classified primarily using the empirical reference distributions derived from laboratory-generated hybrids.

### Validation of hybrid assignments using ADMIXTURE, principal component (PCA), and NeighborNet analyses

To assess the robustness of HIest-based hybrid assignments, supervised ADMIXTURE, PCA, and NeighborNet analyses were performed as complementary genomic analyses. For supervised ADMIXTURE analysis, diagnostic SNPs were selected using the core filtering criteria described above for the HIest analysis, followed by additional filtering to construct a robust diagnostic SNP panel for the combined laboratory and field dataset. The resulting panel comprised 51 diagnostic SNPs. Supervised ADMIXTURE analysis was performed for 436 individuals at K = 2 using the parental samples as predefined reference populations; for graphical presentation, 25 individuals with > 50% missing genotypes across the final 51-SNP panel were omitted from the plot, leaving 411 individuals (122 laboratory samples, 120 field samples from 2023, and 169 field samples from 2024) in the ADMIXTURE plot.

PCA and NeighborNet analysis were performed using a more extensive, independently filtered genome-wide SNP dataset comprising 2,756 SNPs from 416 individuals (122 laboratory samples; 125 and 169 field samples from 2023 and 2024, respectively). PCA was performed using glPca in adegenet, and NeighborNet networks were constructed using SplitsTree v6.6.1. These analyses assessed the genomic placement of field-collected individuals relative to parental and laboratory-generated hybrid classes. Detailed SNP filtering criteria and analytical procedures for ADMIXTURE, PCA, and NeighborNet analyses are provided in the Supporting Information.

### Comparison with single-locus hybrid detection

To compare the sensitivity of genome-wide MIG-seq with that of traditional single-locus approaches, we amplified DNA from all MIG-seq-identified hybrids using hybrid diagnostic primers developed by Li et al. (Li et al., 2021). Due to the low amount of DNA remaining, we used Tks Gflex polymerase (Takara Bio, Kusatsu, Japan). PCR reactions (total volume, 25 µL) contained: 0.5 µL genomic DNA template, 0.5 µL Tks Gflex DNA Polymerase (1.25 U/ µL; Takara Bio), 12.5 µL 2× Gflex PCR Buffer (Takara Bio), 0.5 µL of two forward primers and reverse primer (0.2 µM each), and 10.5 µL nuclease-free water. The thermal cycling conditions of the ProFlex PCR System (Applied Biosystems, Foster City, CA, USA) were as follows: initial denaturation at 94°C for 1 min; 30 cycles of 98°C for 10 s, primer-specific annealing at 55°C for 15 s, and 68°C for 90 s. PCR banding patterns were compared with genome-wide hybrid assignments to evaluate the ability of the single-locus marker to detect advanced-generation hybrids.

### Pre-mating reproductive isolation: chemotaxis assay

Pre-mating reproductive isolation was assessed using a chemotaxis assay modified from Shinya et al. (2015). Approximately 100 test adult nematodes were placed at the center of a 90-mm agar plate, and volatile cues from 10 virgin individuals of the opposite sex were presented against a water control at opposite sides of the plate. Sodium azide was used to immobilize responding nematodes in the cue and control zones. Assays were conducted for 6 h in darkness. To measure the preference toward the test chemical cue, a chemical index (CI) was calculated as: ([Number of nematodes in the cue zone] – [Number of nematodes in the control zone])/([Number of nematodes in the test cue zone] + [Number of nematodes in the control zone]).

We tested conspecific interactions within *B. xylophilus* T4 and reciprocal heterospecific interactions between *B. xylophilus* T4 and either *B. m. mucronatus* Un-1 or *B. m. kolymensis* TCS00 (Table S1). Both male responses to females and female responses to males were examined. Each combination was replicated 3–7 times. Differences in CIs among treatments were assessed using one-way analysis of variance (ANOVA) followed by the Tukey-Kramer test. Detailed assay conditions are provided in the Supporting Information.

### Post-mating reproductive isolation: fecundity and hatchability

Post-mating reproductive isolation was assessed by measuring fecundity and egg hatchability in conspecific and interspecific crosses, following Shinya et al. (2015). Each virgin female was paired with three virgin males to ensure adequate sperm transfer. Conspecific crosses were performed for *B. xylophilus* T4, *B. m. mucronatus* Un-1, and *B. m. kolymensis* TCS00, together with reciprocal interspecific crosses between *B. xylophilus* and each of the two *B. mucronatus* subspecies (Table S4). Each crossing combination was replicated 11–23 times using independently isolated individuals.

Mated females were transferred daily to fresh plates until oviposition ceased. Fecundity was calculated as the total number of eggs laid by each female over the reproductive period. Egg hatchability was assessed at 24 h after oviposition and calculated as the proportion of eggs that hatched. Trials in which females escaped during the assay were excluded to exclude the possibility that the male and female did not mate. Females that produced no eggs were excluded from both fecundity and hatchability analyses; females producing at least one egg were included in the hatchability analysis. Differences in fecundity and hatchability among crossing combinations were evaluated using one-way ANOVA followed by the Tukey-Kramer test. Detailed experimental procedures are provided in the Supporting Information.

## Results

### Empirical reference distributions improve HIest-based hybrid classification

To evaluate whether hybrid generations could be distinguished using genome-wide SNP data, we examined the distribution of laboratory-generated individuals of known pedigree in S–H space, defined by the HIest hybrid index (S, the proportion of *B. xylophilus* ancestry) and interspecific heterozygosity (H) (Figure 1). Parental, F1, B1, and B2 classes formed distinct distributions in both species comparisons, demonstrating that MIG-seq-derived diagnostic SNPs provided sufficient resolution to distinguish major hybrid classes.

**Figure 1.**
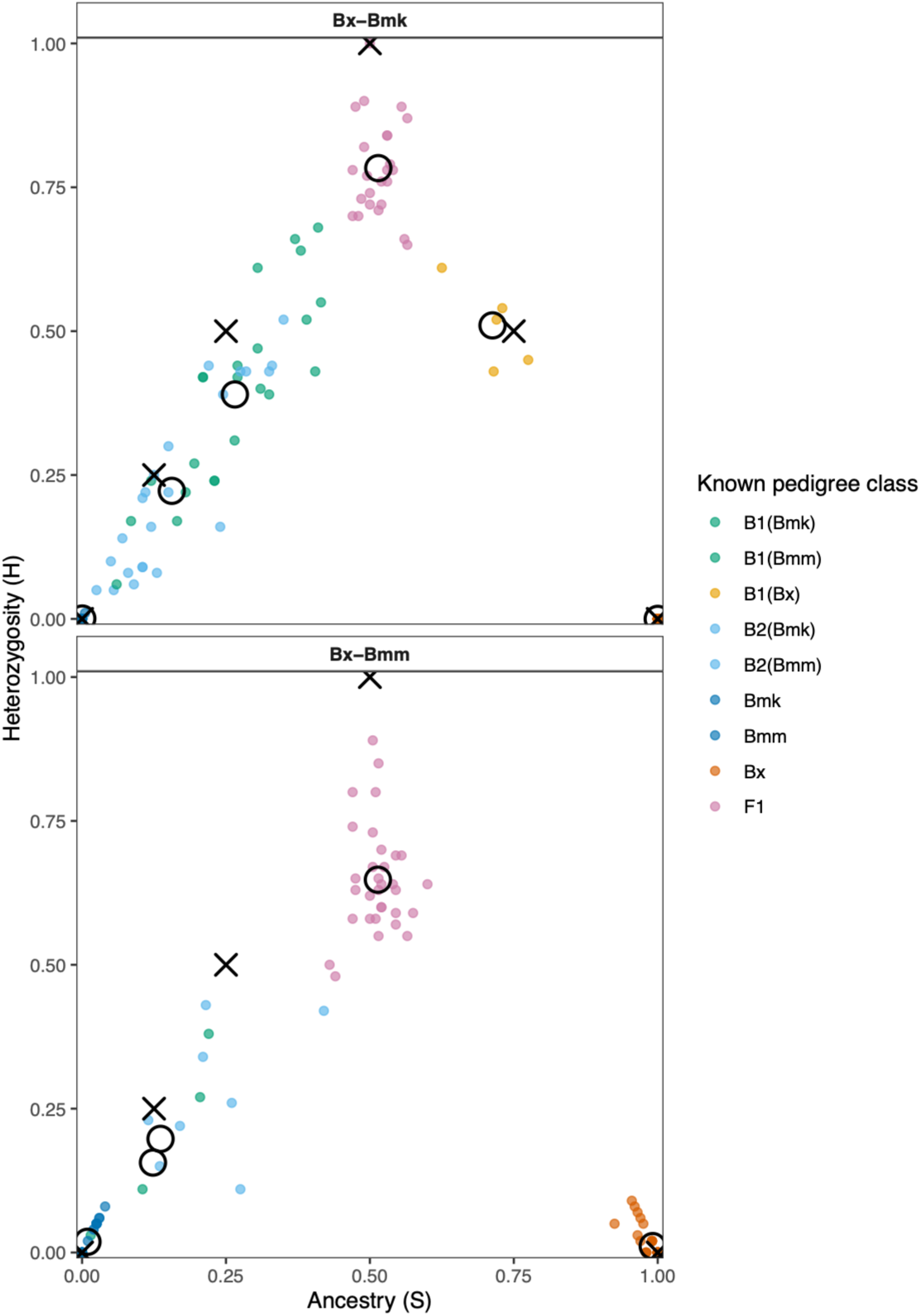
Theoretical and empirical positions of hybrid classes in ancestry (S)– heterozygosity (H) space. S and H were estimated by HIest for laboratory individuals of known hybrid class in the *Bursaphelenchus xylophilus*–*Bursaphelenchus mucronatus kolymensis* (top) and *B. xylophilus*–*B. mucronatus mucronatus* (bottom) datasets. Dots represent individual samples. Crosses indicate theoretical class positions, and large open circles indicate empirical class centers calculated from individuals of known pedigree. In both datasets, hybrid classes formed well-separated distributions in S–H space, demonstrating that multiplexed inter-simple sequence repeat genotyping by sequencing (MIG-seq)-derived single-nucleotide polymorphism (SNP) data combined with HIest analysis provided sufficient resolution to distinguish major hybrid classes. Empirical heterozygosity was generally lower than theoretical expectations, with pronounced downward shifts in F1 individuals. Theoretical and empirical values for each class are provided in Table 1.

**Table 1.** Theoretical S and H values were compared with empirical estimates obtained from laboratory-generated individuals of known pedigree for the *B. xylophilus*–*B. m. kolymensis* and *B. xylophilus*–*B. m. mucronatus* datasets. Empirical values are presented as means ± SD; these were used as class-specific references for the assignment of field-collected individuals. Across both datasets, empirical heterozygosity showed a consistent downward shift relative to theoretical expectations; the largest differences were observed in F1 individuals. Labels in parentheses indicate the recurrent parental taxon in backcross classes.

| Dataset | Class | n | S_theoretical | H_theoretical | S_empirical | H_empirical | SD_S | SD_H | Distance |
| --- | --- | --- | --- | --- | --- | --- | --- | --- | --- |
| Bx-Bmk | Bmk | 26 | 0 | 0 | 0.027 | 0.026 | 0.042 | 0.037 | 0.037 |
| Bx-Bmk | B2(Bmk) | 16 | 0.125 | 0.25 | 0.159 | 0.22 | 0.059 | 0.05 | 0.045 |
| Bx-Bmk | B1(Bmk) | 22 | 0.25 | 0.5 | 0.322 | 0.492 | 0.072 | 0.102 | 0.072 |
| Bx-Bmk | F1 | 21 | 0.5 | 1 | 0.512 | 0.8 | 0.026 | 0.077 | 0.201 |
| Bx-Bmk | B1(Bx) | 7 | 0.75 | 0.5 | 0.67 | 0.551 | 0.086 | 0.092 | 0.095 |
| Bx-Bmk | Bx | 16 | 1 | 0 | 1 | 0 | 0 | 0 | 0 |
| Bx-Bmm | Bmm | 78 | 0 | 0 | 0.041 | 0.032 | 0.051 | 0.041 | 0.052 |
| Bx-Bmm | B2(Bmm) | 65 | 0.125 | 0.25 | 0.182 | 0.198 | 0.062 | 0.057 | 0.078 |
| Bx-Bmm | B1(Bmm) | 83 | 0.25 | 0.5 | 0.319 | 0.452 | 0.074 | 0.088 | 0.083 |
| Bx-Bmm | F1 | 91 | 0.5 | 1 | 0.508 | 0.676 | 0.031 | 0.081 | 0.324 |
| Bx-Bmm | B1(Bx) | 74 | 0.75 | 0.5 | 0.691 | 0.447 | 0.069 | 0.084 | 0.084 |
| Bx-Bmm | Bx | 82 | 1 | 0 | 0.982 | 0.021 | 0.028 | 0.03 | 0.027 |

However, empirical class distributions differed systematically from theoretical Mendelian expectations (Figure 1; Table 1). In particular, observed interspecific heterozygosity was consistently lower than expected, leading to downward displacement of several hybrid classes in S–H space. This deviation was most pronounced in F1 individuals and was greater in the *B. xylophilus*–*B. m. mucronatus* dataset than in the *B. xylophilus*–*B. m. kolymensis* dataset.

Classification based on empirical class centers substantially outperformed classification based on theoretical expectations (Figure 2). Accuracy increased from 63.2% to 77.6% for the *B. xylophilus*–*B. m. kolymensis* dataset and from 63.2% to 81.1% for the *B. xylophilus*–*B. m. mucronatus* dataset. Therefore, we used empirical reference distributions derived from pedigree-confirmed laboratory individuals to classify field-collected samples. Detailed class-specific S and H distributions and classification outcomes are provided in Table S8.

**Figure 2.**
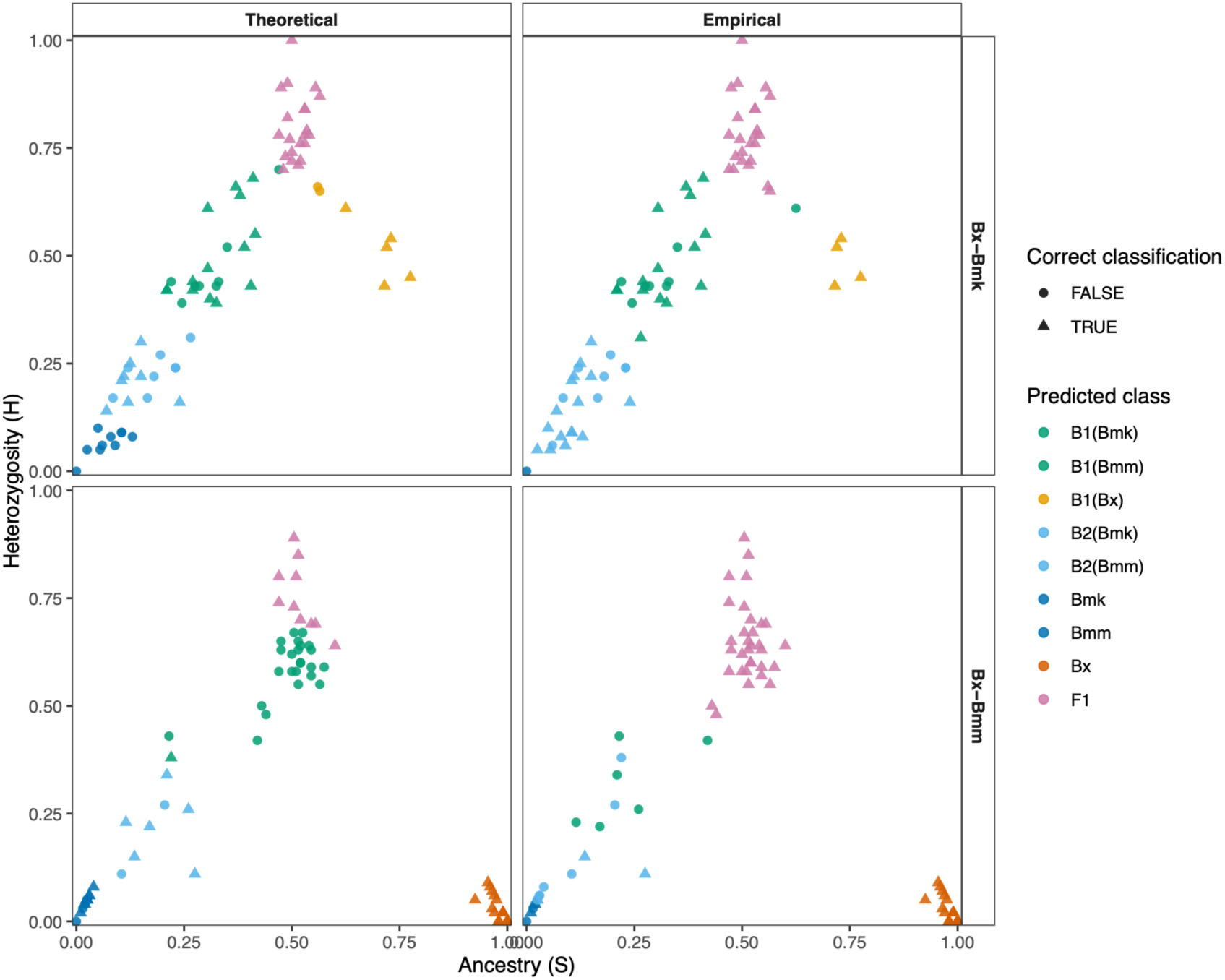
Classification accuracy of theoretical and empirical approaches in assigning individuals to hybrid classes based on their positions in S–H space. Predictions were evaluated against known pedigrees of laboratory-generated individuals that were independent of parental reference individuals used for HIest estimation. For the empirical approach, classification performance was evaluated using leave-one-out cross-validation (LOOCV). Both approaches recovered the major hybrid categories, although classification performance differed among pedigree classes. The empirical approach, which used class-specific centers and variation derived from observed data (Table 1), improved overall classification accuracy relative to the theoretical approach under observed marker and missing-data conditions. Detailed overall and class-specific classification accuracy data are provided in Table S8.

### Natural hybrids and advanced-generation introgression were detected in both sampling years

Species-specific PCR screening showed that *B. xylophilus* and *B. mucronatus* frequently co-occurred within individual vector beetles. Both species were detected in 31 of 115 beetle-derived populations collected in 2023 (27.0%) and in 20 of 126 populations collected in 2024 (15.9%) (Figure S1; Table S6).

Application of the empirical HIest framework to field-collected individuals identified nine high-confidence hybrids: three collected in 2023 and six collected in 2024 (Figure 3; Table 2). The 2023 hybrids were 4-g, 4-i, and 14-h, and the 2024 hybrids were 3-f, 5-r, 5-t, 12-k, 15-f, and 15-i. These individuals originated from six different *M. alternatus* beetles, demonstrating recurrent hybrid formation across independent beetle-derived populations and consecutive years.

**Figure 3.**
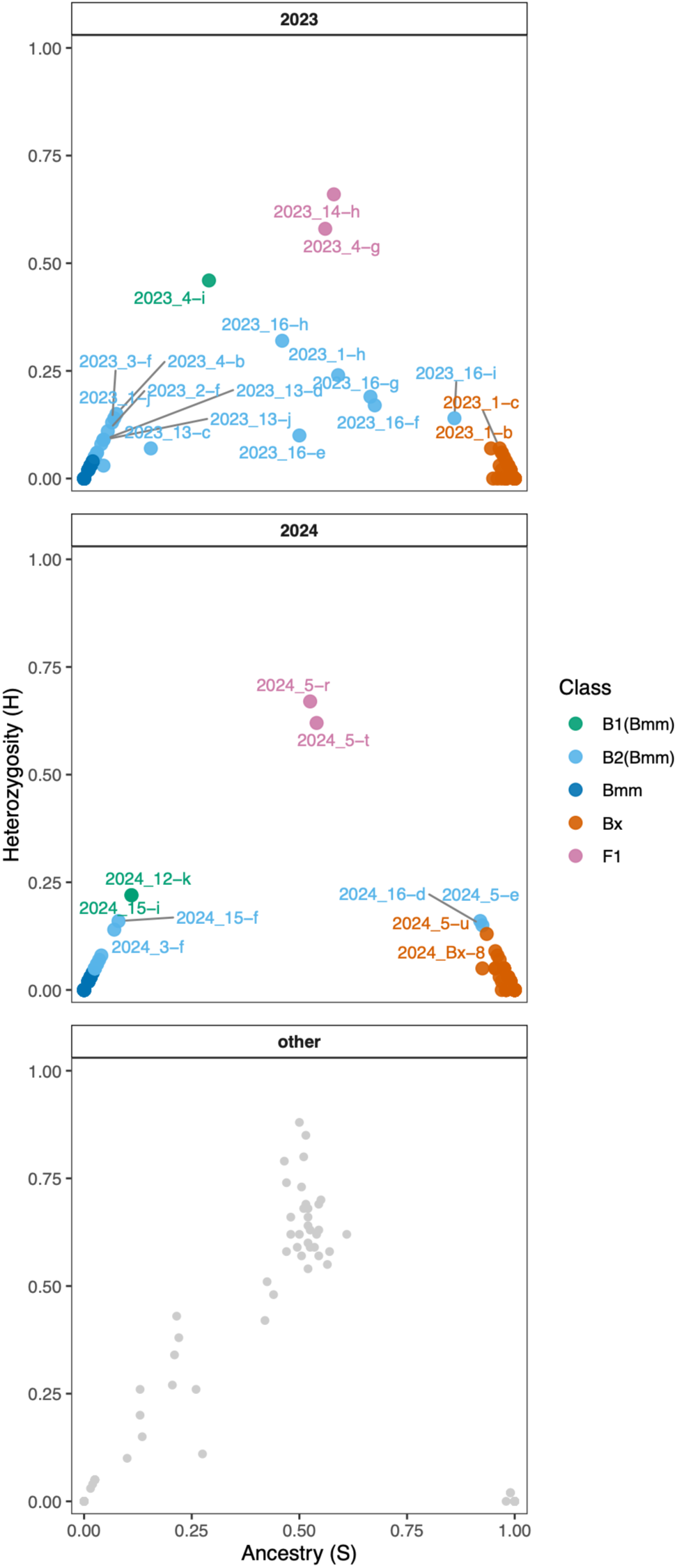
Empirical assignment of hybrid classes in field samples. Hybrid classes of field-collected individuals sampled in 2023 and 2024 were assigned based on their positions in S–H space using an empirical reference approach. Class assignments were performed using standardized Euclidean distances to class-specific empirical centers and standard deviations (SDs) derived from laboratory individuals of known pedigree (Table 1). Class assignments were restricted to classes represented in the reference dataset. However, some individuals occupied intermediate positions in S–H space that did not closely correspond to the empirical centers of the represented reference classes, illustrating the continuous nature of hybrid ancestry. Grey dots represent individuals from the laboratory reference dataset, shown for context but not reclassified. Colored dots indicate field-collected individuals, with colors corresponding to the nearest empirical classes. Panels are separated by sampling year, selected field-collected individuals are labeled, including the nine high-confidence field hybrids. Individual ancestry estimates and final assignments are provided in Table S9.

**Table 2.** Genomic ancestry and single-locus genotypes of high-confidence field hybrids detected among field-collected nematodes in 2023 and 2024. Primary empirical assignment was based on class-specific standardized distances to empirical reference class centers, with the second-closest class and distance margin shown to indicate classification separation. Ordinary Euclidean (OE) assignment using the same empirical class centers without class-specific standardization is provided as a sensitivity check. The supervised ADMIXTURE ancestry coefficient for *B. xylophilus* [q(Bx)], overall genomic interpretation, and single-locus hybrid diagnostic assay results are shown. Overall genomic interpretations were based on the combined evidence from HIest and ADMIXTURE analysis. The terms F1-like and advanced-backcross-like describe genomic ancestry patterns and should not be interpreted as exact pedigrees. Bx, *B. xylophilus*; Bmm, *B. m. mucronatus*.

| Sample | Year | n SNP (Hlest) | S | H | Empirical assignment | Second class | Distance margin | OE sensitivity check | q(Bx) | Interpretation category | Interpretation | Single-locus hybrid-diagnostic assay |
| --- | --- | --- | --- | --- | --- | --- | --- | --- | --- | --- | --- | --- |
| 14-h | 2023 | 29 | 0.58 | 0.66 | F1 | B2(Bmm) | 3.296 | F1 | 0.56 | Strong F1-like | Strong F1-like signal; both methods agree and ADMIXTURE is intermediate. | hybrid |
| 4-g | 2023 | 26 | 0.56 | 0.58 | F1 | B2(Bmm) | 3.153 | F1 | 0.54 | Strong F1-like | Strong F1-like signal; both methods agree and ADMIXTURE is intermediate. | hybrid |
| 4-i | 2023 | 34 | 0.29 | 0.46 | B1(Bmm) | B2(Bmm) | 0.196 | F1 | 0.22 | Hybrid, generation uncertain | Hybrid signal supported by both methods; F1 vs backcross generation is ambiguous, especially under ordinary distance. | hybrid |
| 5-r | 2024 | 32 | 0.53 | 0.67 | F1 | B2(Bmm) | 4.451 | F1 | 0.53 | Strong F1-like | Strong F1-like signal; both methods agree and ADMIXTURE is intermediate. | hybrid |
| 5-t | 2024 | 39 | 0.54 | 0.62 | F1 | B2(Bmm) | 3.899 | F1 | 0.53 | Strong F1-like | Strong F1-like signal; both methods agree and ADMIXTURE is intermediate. | hybrid |
| 12-k | 2024 | 37 | 0.11 | 0.22 | B1(Bmm) | B2(Bmm) | 0.149 | B1(Bmm) | 0.07 | Backcross-like | Backcross-like toward Bmm; both methods agree, but B1 vs B2 is only weakly separated. | Bm |
| 15-f | 2024 | 38 | 0.08 | 0.16 | B2(Bmm) | B1(Bmm) | 0.287 | B2(Bmm) | 0.04 | Backcross-like | Backcross-like toward Bmm; both methods agree on a hybrid class, but B1/B2 resolution is limited. | Bm |
| 15-i | 2024 | 37 | 0.11 | 0.22 | B1(Bmm) | B2(Bmm) | 0.149 | B1(Bmm) | 0.06 | Backcross-like | Backcross-like toward Bmm; both methods agree, but B1 vs B2 is only weakly separated. | Bm |
| 3-f | 2024 | 39 | 0.07 | 0.14 | B2(Bmm) | B1(Bmm) | 0.341 | B2(Bmm) | 0.02 | Backcross-like | Backcross-like toward Bmm; both methods agree on a hybrid class, but B1/B2 resolution is limited. | Bm |

Individuals 4-g, 14-h, 5-r, and 5-t showed relatively balanced genomic ancestry and occupied positions consistent with F1-like or other early-generation hybrid ancestry. In contrast, individuals 4-i, 3-f, 12-k, 15-f, and 15-i showed strongly asymmetric ancestry patterns consistent with advanced-generation backcrossing. Three hybrid pairs were recovered from the same beetle-derived populations: 4-g and 4-i from beetle no. 52 in 2023, 5-r and 5-t from beetle no. 68 in 2024, and 15-f and 15-i from beetle no. 121 in 2024.

Additional field-collected individuals occupied positions close to hybrid reference classes in S–H space but were not included in the high-confidence set because their assignments were not consistently supported across the complementary genomic analyses. Results for these HIest-only candidate hybrids are reported separately in Table S9.

### ADMIXTURE and NeighborNet analyses supported field hybrid assignments

Supervised ADMIXTURE analysis provided ancestry estimates that were broadly concordant with the HIest-based assignments (Figure 4). Laboratory parental individuals showed ancestry coefficients close to the corresponding parental extremes, whereas pedigree-confirmed F1 and backcross individuals showed intermediate or asymmetric ancestry proportions consistent with their known pedigrees.

**Figure 4.**
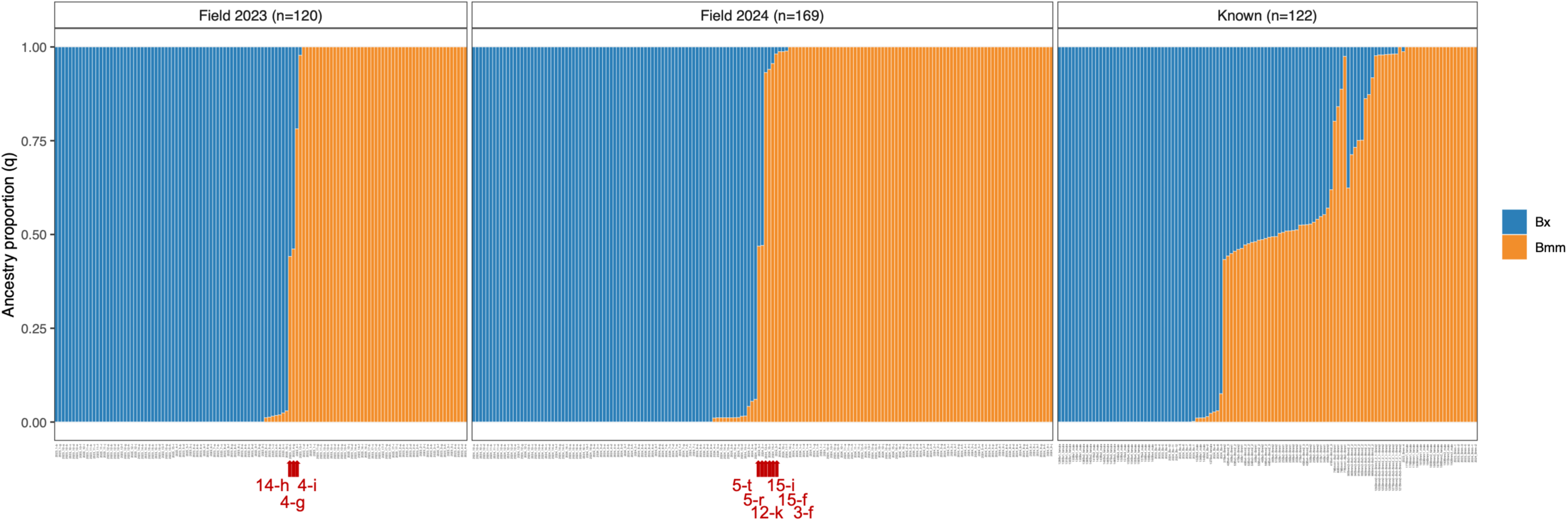
Supervised ADMIXTURE analysis (K = 2) of laboratory and field samples was conducted using 51 diagnostic SNPs derived from MIG-seq data. Parental samples of *B. xylophilus* (Bx) and *B. m. mucronatus* (Bmm) were used as predefined reference populations. For graphical presentation, 25 individuals with > 50% missing genotypes across the final 51-SNP panel were omitted, leaving a total of 411 individuals. The dataset comprised field-collected individuals sampled in 2023 and 2024, together with laboratory reference individuals. The Known panel includes pure parental reference strains of Bx and Bmm, as well as laboratory-generated F1, B1, and B2 individuals used to anchor ancestry estimation. Field-collected individuals were analyzed without predefined ancestry assignments. Each vertical bar represents one individual; colors indicate estimated ancestry proportions (*q*), with blue and orange corresponding to Bx and Bmm ancestry, respectively. Most field-collected individuals showed ancestry proportions close to one of the parental species; a few showed intermediate or strongly asymmetric ancestry coefficients overlapping those observed in laboratory-defined hybrid classes. Red arrows indicate nine high-confidence field hybrids (4-g, 4-i, 14-h, 3-f, 5-r, 5-t, 12-k, 15-f, 15-i). Ancestry proportions inferred by supervised ADMIXTURE analysis were interpreted comparatively relative to laboratory individuals of known pedigree, rather than as definitive hybrid classifications.

Among field-collected individuals, those classified as F1-like by HIest showed relatively balanced ancestry proportions, whereas the putative advanced backcrosses showed strongly asymmetric ancestry. Thus, the four F1-like individuals were distinguishable from the strongly parentally biased ancestry profiles of individuals 4-i, 3-f, 12-k, 15-f, and 15-i. Individual ancestry coefficients are summarized in Table 2 and provided in full in Table S9.

NeighborNet analysis based on a broader dataset of 2,756 genome-wide SNPs provided complementary support for these assignments (Figure 5). The parental taxa occupied differentiated regions of the network, while pedigree-confirmed laboratory hybrids were located between the parental groups or closer to the recurrent parental taxon according to their pedigrees. Field individuals with relatively balanced ancestry generally occupied intermediate genomic positions, whereas putative advanced backcrosses were positioned closer to one of the parental groups.

**Figure 5.**
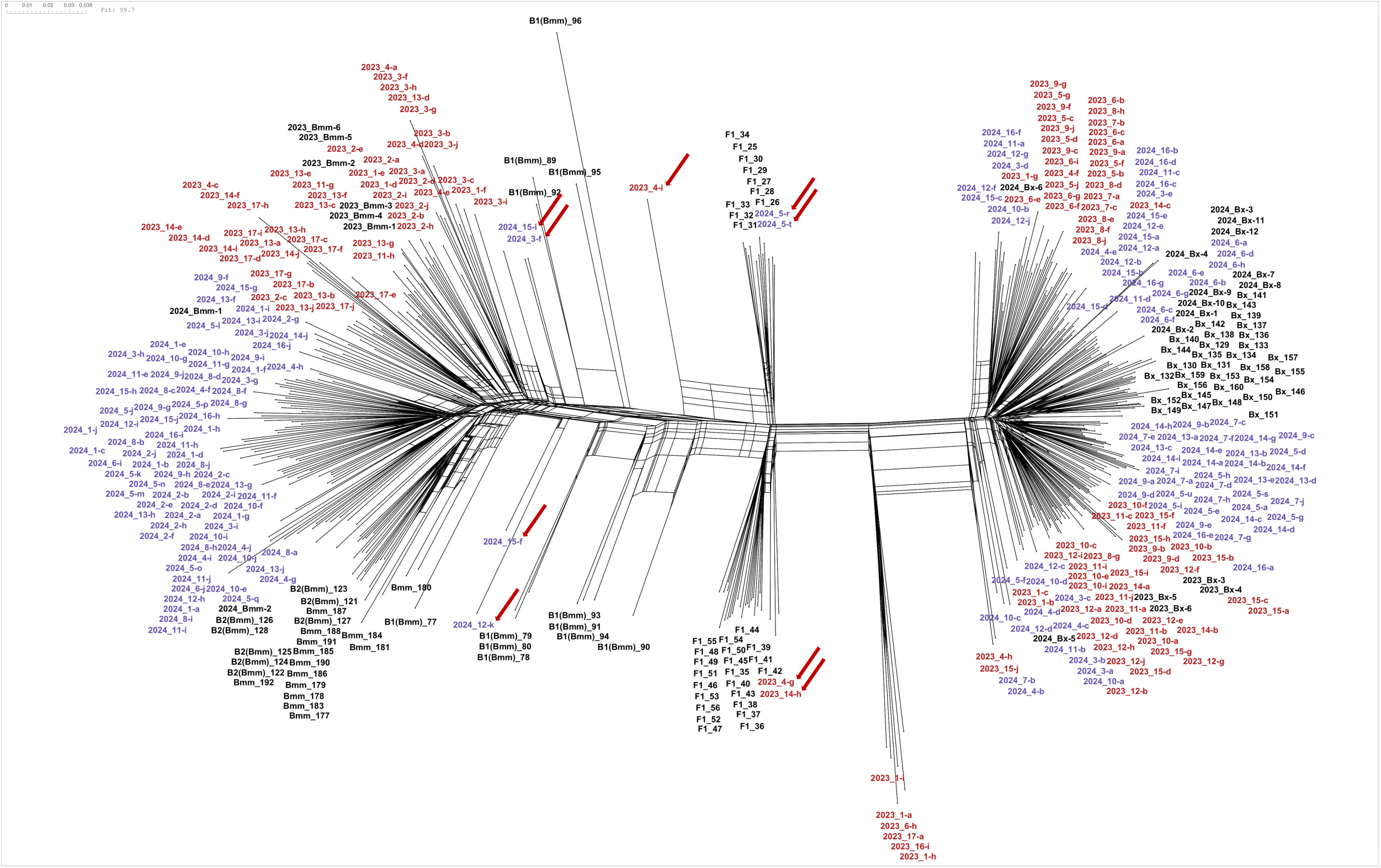
NeighborNet network analysis based on genome-wide SNP data. The network was constructed using 2,756 independently filtered genome-wide SNPs from 416 laboratory-generated and field-collected samples. Distances were calculated using uncorrected p-distances, and ambiguous sites were handled using the MatchStates option. Laboratory samples, including parental and known-pedigree hybrid individuals, are indicated in black, whereas field samples are colored by year (red, 2023; blue, 2024). Two major groups corresponding to *B. xylophilus* and *B. mucronatus* are connected by a reticulate structure, with relatively balanced hybrids occupying intermediate positions and parentally biased individuals positioned closer to one of the parental groups. The nine high-confidence field hybrids are labeled.

PCA based on the same broader genome-wide SNP dataset produced a generally concordant pattern (Figure S2).

### Single-locus markers failed to detect advanced backcrosses

All nine high-confidence field hybrids identified by genome-wide analyses were re-genotyped using the single-locus hybrid-diagnostic marker developed by Li et al. (2021). Five individuals (4-g, 4-i, 14-h, 5-r, and 5-t) displayed the dual-banding pattern expected for hybrids and were therefore detectable using the single-locus assay (Figure 6; Table 2).

**Figure 6.**
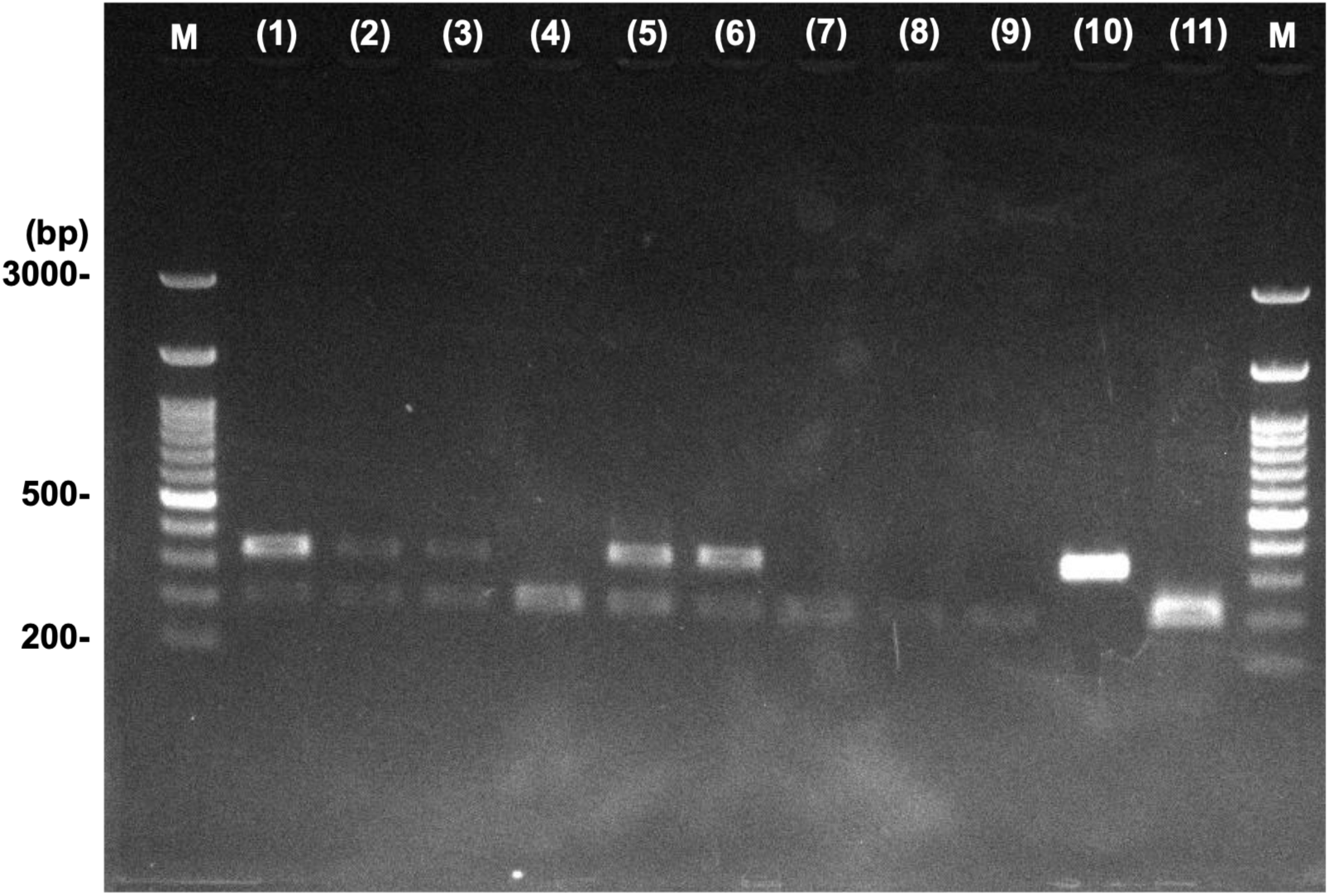
DNA banding patterns obtained using single-locus hybrid diagnostic primers. DNA banding patterns obtained using the single-locus hybrid-diagnostic primers developed by Li et al. (2021) are shown for field-collected individuals examined in genome-wide SNP analyses. Lanes (1)–(9) correspond to field-collected individuals: (1) 4-g, (2) 4-i, (3) 14-h, (4) 3-f, (5) 5-r, (6) 5-t, (7) 12-k, (8) 15-f, and (9) 15-i. As pure parental controls, lane 10 contains *B. xylophilus* Ka4 and lane 11 contains *B. m. mucronatus* Ioujima. Lane M contains a molecular size marker (100–3,000 bp). Individuals 4-g, 4-i, 14-h, 5-r, and 5-t showed the dual-banding pattern expected for hybrids, whereas 3-f, 12-k, 15-f, and 15-i showed only the *B. m. mucronatus*-type band.

In contrast, individuals 3-f, 12-k, 15-f, and 15-i produced only the *B. mucronatus*-type band and were indistinguishable from pure *B. mucronatus* using the single-locus marker. Nevertheless, these individuals carried alleles from both parental taxa at multiple genome-wide diagnostic SNPs and showed strongly asymmetric ancestry consistent with advanced backcrossing.

Thus, the single-locus assay failed to detect four of the nine hybrids identified by genome-wide analyses (44.4%). Detection failure was specifically associated with individuals showing the most strongly parentally biased genomic ancestry, demonstrating that single-locus markers can fail to identify advanced-generation introgression.

### No evidence of pre-mating discrimination between species

Chemotaxis responses did not differ significantly between conspecific and heterospecific cue combinations in either species comparison or response direction (Figure 7; Table S4). Male responses to female cues were not significantly different among treatments for either the *B. xylophilus*–*B. m. mucronatus* comparison (F_3, 15_ = 1.182, *P* = 0.3498) or the *B. xylophilus*–*B. m. kolymensis* comparison (F_3, 17_ = 1.684, *P* = 0.2082).

**Figure 7.**
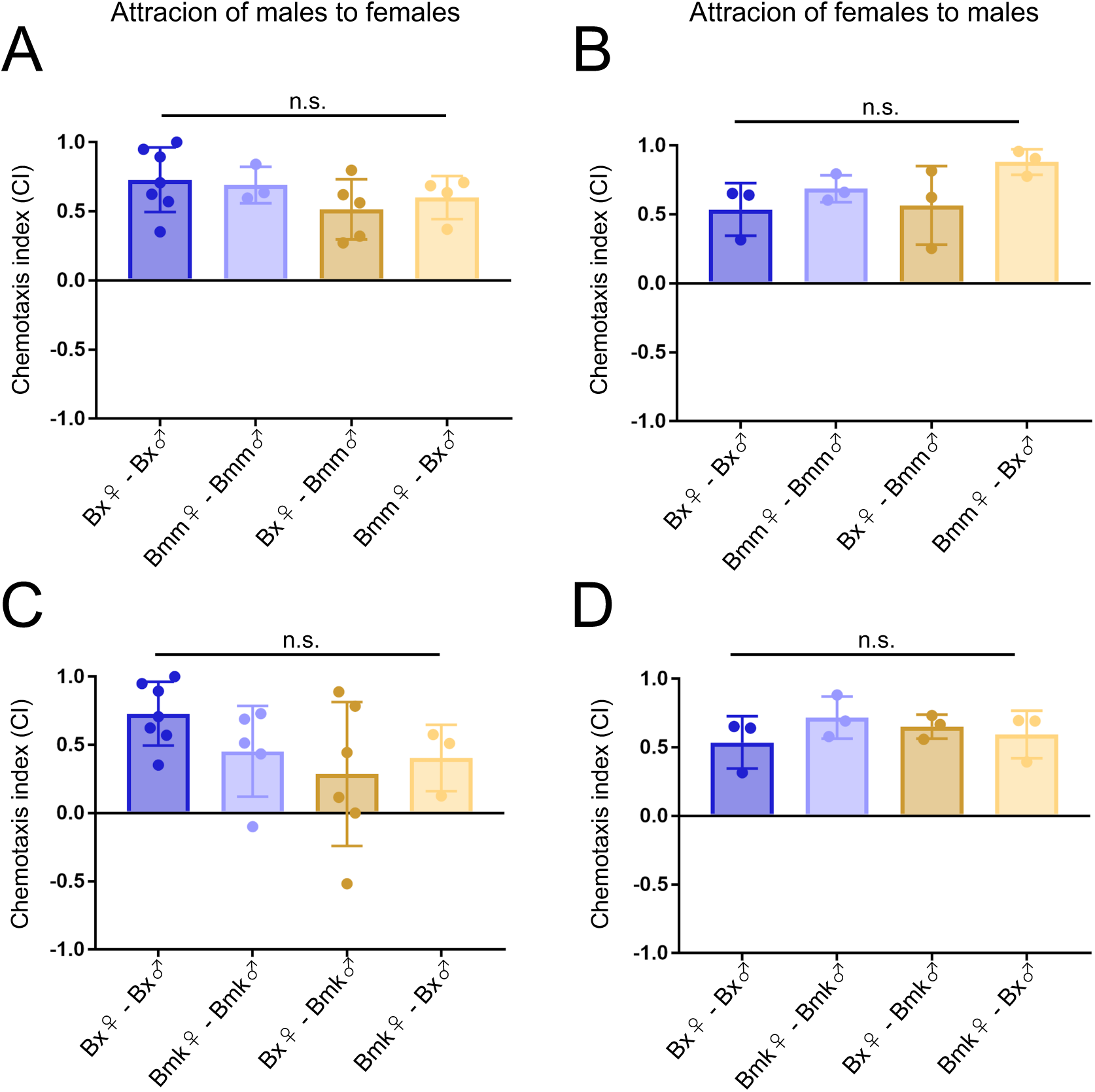
Chemotactic attraction to sex-specific chemical cues. Chemotaxis assays measured attraction to chemical cues produced by the opposite sex. In each panel, the x-axis indicates the sex of the odor source, with females shown on the left and males on the right, regardless of species. Attraction was measured toward sex-specific chemical cues rather than toward live individuals. (A, B) Assays between *B. xylophilus* (Bx) and *B. m. mucronatus* (Bmm); (C, D) assays between Bx and *B. m. kolymensis* (Bmk). Bx strain T4, Bmm strain Un-1, and Bmk strain TCS00 were used. No significant differences were detected among conspecific and heterospecific cue combinations. Replication numbers are provided in Table S4.

Female responses to male cues were also indistinguishable among conspecific and heterospecific treatments for both the *B. xylophilus*–*B. m. mucronatus* comparison (F_3, 8_ = 2.158, *P* = 0.1711) and the *B. xylophilus*–*B. m. kolymensis* comparison (F_3, 8_ = 0.7445, *P* = 0.5551). These results provide no evidence of pre-mating isolation through discrimination of opposite-sex volatile cues.

### Post-mating reproductive barriers were asymmetric and cross-dependent

Post-mating reproductive performance varied among crossing combinations (Figure 8; Table S5). Egg-laying success was 65.2-92.3% among conspecific crosses and 66.7-91.3% among interspecific crosses. Trials in which no egg-laying was observed were excluded from analysis because mating might not have occurred.

**Figure 8.**
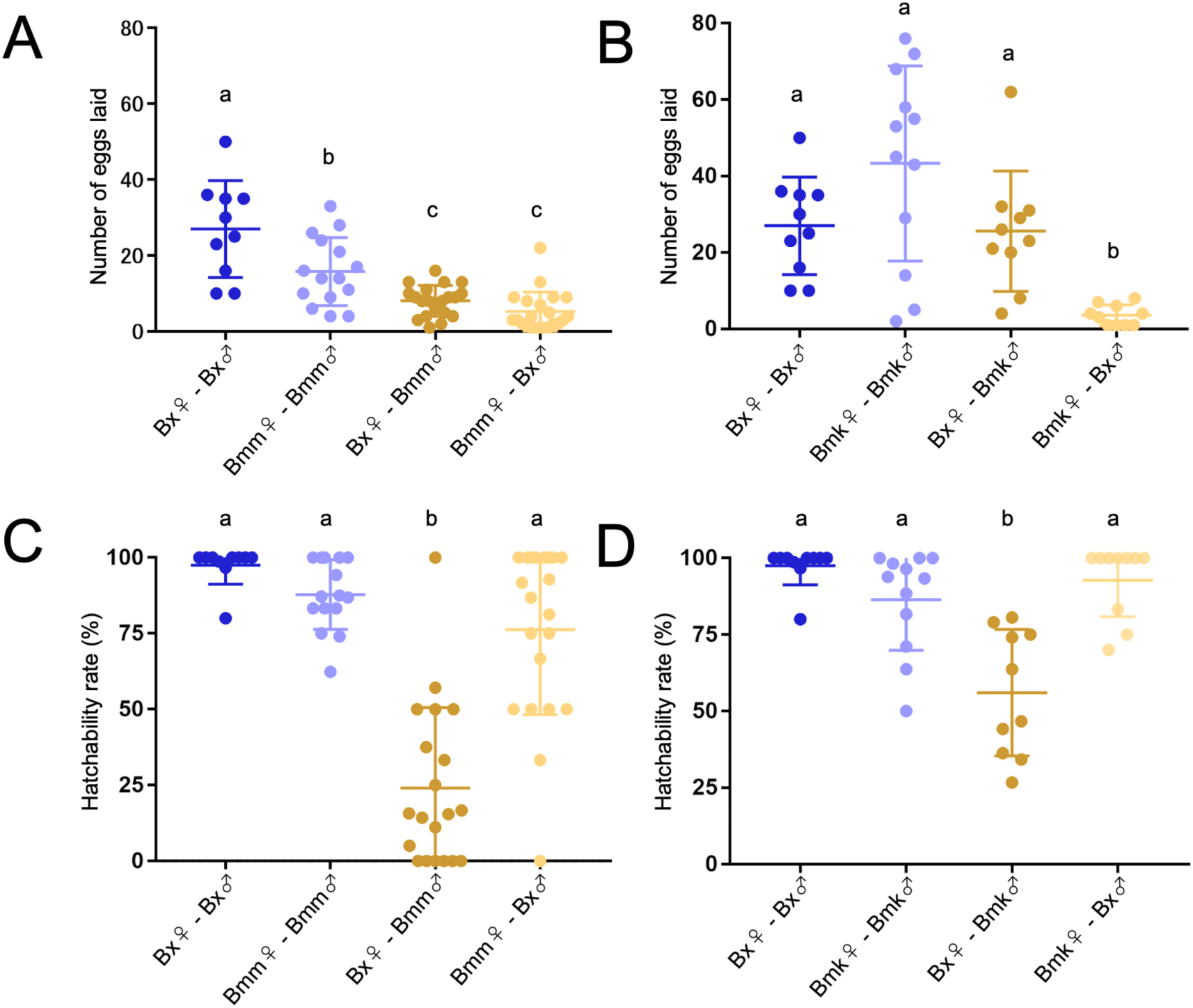
Fecundity and egg hatchability in intra- and interspecific crosses. Numbers of eggs laid by egg-producing females (A, B) and 24-h hatchability rates (C, D) are shown for crosses among *B. xylophilus* (Bx), *B. m. mucronatus* (Bmm), and *B. m. kolymensis* (Bmk). In each panel, the x-axis indicates the sex of the focal individual, with females shown on the left and males on the right, regardless of species. (A, C) Crosses between Bx and Bmm; (B, D) crosses between Bx and Bmk. *Bx* strain T4, Bmm strain Un-1, and Bmk strain TCS00 were used. Crosses producing no eggs were included in the analysis of egg-laying success but not in conditional analyses of positive fecundity and hatchability. Different letters indicate significant pairwise differences after adjustment for multiple comparisons. Complete trial numbers, descriptive statistics, model results, and pairwise comparisons are provided in Table S5.

Among females that produced eggs, mean fecundity was 27.0 ± 4.04 (mean ± standard error [SE]) eggs for *B. xylophilus*, 15.8 ± 2.32 eggs for *B. m. mucronatus*, and 43.3 ± 7.38 eggs for *B. m. kolymensis* conspecific crosses. Among interspecific crosses, mean fecundity was lowest when *B. m. kolymensis* females were crossed with *B. xylophilus* males (3.6 ± 0.85 eggs).

Hatchability showed a pronounced directional pattern. Crosses with *B. xylophilus* as the maternal parent had reduced hatchability when males were either *B. m. mucronatus* (24.1 ± 26.6%, mean ± SD) or *B. m. kolymensis* (56.0 ± 20.7%). In contrast, hatchability was substantially higher in the reciprocal crosses: 76.3 ± 28.0% for *B. m. mucronatus* females crossed with *B. xylophilus* males and 92.8 ± 12.0% for *B. m. kolymensis* females crossed with *B. xylophilus* males.

All four reciprocal interspecific crossing directions produced at least some viable offspring. Thus, reproductive isolation was incomplete, although post-mating barriers were strongly dependent on cross direction and the *B. mucronatus* subspecies involved.

## Discussion

This study provides the first genome-wide evidence for ongoing natural hybridization between the invasive pinewood nematode *B. xylophilus* and the native *B. mucronatus* in Japan. We identified nine high-confidence hybrids among 342 field-collected nematodes, including three individuals sampled in 2023 and six sampled in 2024 (Figure 3; Table 2). These individuals originated from six independent *M. alternatus* beetles and exhibited genomic patterns ranging from relatively balanced, early-generation ancestry to strongly asymmetric, advanced-backcross-like ancestry. Their occurrence across multiple beetles and in consecutive years indicates that hybridization is recurrent rather than attributable to a single mating event or beetle-derived population. Moreover, the presence of backcross-like individuals in both sampling years provides strong evidence that hybridization has progressed beyond initial F1 formation and is contributing to multigenerational introgression. This result extends the findings of Li et al. (2021), who detected natural hybrids using a single nuclear marker but could not comprehensively characterize later-generation ancestry. The frequent co-occurrence of both species within individual vector beetles (27.0% of beetle-derived populations in 2023 and 15.9% in 2024) provides repeated opportunities for interspecific contact during the phoretic phase of the life cycle (Figure S1; Table S6), consistent with previous observations of sympatry within individual vectors in Japan (Ozawa et al., 2021).

A central methodological contribution of this study is the pedigree-based calibration of hybrid classification in S–H space. Laboratory-generated individuals of known ancestry formed distinguishable parental, F1 and backcross distributions, confirming that HIest estimates derived from MIG-seq diagnostic SNPs can resolve major hybrid classes (Figure 1). However, empirical interspecific heterozygosity was systematically lower than theoretical Mendelian expectations, particularly in F1 individuals (Table 1). Consequently, classification based directly on theoretical S and H values led to substantial displacement and misclassification of several hybrid classes. Using empirical class centers and class-specific variation increased classification accuracy from 63.2% to 77.6% for the *B. xylophilus*–*B. m. kolymensis* dataset and from 63.2% to 81.1% for the *B. xylophilus*–*B. m. mucronatus* dataset (Figure 2; Table S8). The downward shift in H may reflect a combination of missing genotypes, allelic dropout, uneven sequencing depth, ascertainment of diagnostic loci, and stochastic segregation across a finite marker panel. Regardless of its precise cause, the improvement obtained using empirical references demonstrates that idealized theoretical positions should not be applied uncritically to genomic data with reduced representation. Pedigree-confirmed calibration is particularly important when distinguishing F1 individuals from later-generation backcrosses whose ancestry distributions may overlap.

Complementary genomic analyses strengthened this interpretation. Supervised ADMIXTURE analysis produced ancestry proportions that were broadly concordant with the HIest assignments, with relatively balanced ancestry observed in early-generation hybrids and strongly parental-biased ancestry observed in putative advanced backcrosses (Figure 4). Because ADMIXTURE analysis is based on a closely related diagnostic SNP signal, it is best regarded as a complementary representation of ancestry rather than a fully independent validation of the HIest classifications. By contrast, NeighborNet analysis and PCA used a broader dataset containing 2,756 genome-wide SNPs. NeighborNet analysis placed laboratory-generated and field-collected early-generation hybrids between the parental taxa, while advanced-backcross-like individuals were positioned closer to the inferred recurrent parental group (Figure 5). PCA produced a generally concordant pattern using the same broader dataset (Figure S2). Agreement between the diagnostic SNP analyses and the broader genome-wide analyses reduces the likelihood that the nine high-confidence assignments were artefacts of a small marker panel. Simultaneously, individuals supported only by their position in HIest S–H space were conservatively retained as candidate rather than confirmed hybrids (Table S9).

A comparison of the single-locus assay results illustrates how later-generation introgression can remain cryptic. Four of the nine high-confidence hybrids (3-f, 12-k, 15-f, and 15-i) were indistinguishable from pure *B. mucronatus* based on the nuclear marker developed by Li et al. (2021), despite carrying alleles from both parental taxa at multiple genome-wide loci (Figure 6; Table 2). Thus, the single-locus assay failed to detect 44.4% of the hybrids identified in genome-wide analyses. This result follows directly from Mendelian segregation. Following an F1 backcross to one parental species, a diagnostic locus has an approximately 50% probability of becoming homozygous for the recurrent-parent allele; after a second backcross, this probability increases to approximately 75%. Consequently, an advanced backcross can retain introgressed ancestry elsewhere in the genome while appearing genetically pure at any individual marker. It was not possible to morphologically distinguish between parental species and hybrids of *B. xylophilus* and *B. mucronatus* (Figure S3); similar underestimation of introgression by morphology or sparse marker sets has been reported in other animal and plant systems (Boyer et al., 2008; Muhlfeld et al., 2009; Sloop et al., 2011). Although the diagnostic panel used for HIest was modest in size, its loci were selected for strong parental differentiation, and its ancestry assignments were calibrated using known pedigrees and supported by analyses of 2,756 broader genome-wide SNPs. These results support the use of multilocus approaches such as MIG-seq or appropriately validated targeted SNP panels when monitoring advanced-generation introgression.

The genomic detection of natural hybrids is mechanistically consistent with incomplete reproductive isolation between the two species. Chemotaxis assays provided no evidence that males or females discriminated between conspecific and heterospecific volatile cues (Figure 7). Therefore, volatile sex attraction signals appear unlikely to constitute a strong pre-mating barrier under our experimental conditions. This finding is consistent with previous observations that pheromone responses can be conserved across *Bursaphelenchus* species, including between *B. xylophilus* and the more distantly related *Bursaphelenchus okinawaensis* (Shinya et al., 2015). Nevertheless, the absence of differential chemotaxis does not exclude other forms of pre-mating isolation in nature, including temporal differences in development, spatial segregation within host trees or vectors, contact-dependent mate recognition, and competitive interactions among males.

Post-mating isolation was more substantial but varied among cross directions and reproductive components (Figure 8; Table S5). Crosses using *B. xylophilus* as the maternal parent showed markedly reduced hatchability when males were either *B. m. mucronatus* (24.1%) or *B. m. kolymensis* (56.0%). In contrast, the reciprocal crosses showed higher hatchability, reaching 76.3% for *B. m. mucronatus* females crossed with *B. xylophilus* males and 92.8% for *B. m. kolymensis* females crossed with *B. xylophilus* males. However, high hatchability did not always correspond to high total reproductive output; *B. m. kolymensis* females crossed with *B. xylophilus* males showed high egg hatchability but markedly reduced fecundity. Therefore, reproductive compatibility cannot be summarized based on hatchability alone and depends on the sequential contributions of mating success, egg production and embryonic survival.

Directional reduction in hatchability is compatible with cytonuclear incompatibility, maternal-effect incompatibility or other asymmetric interactions during embryogenesis, although the present experiments did not distinguish among these mechanisms. In many hybrid systems, mismatches between maternally inherited mitochondrial genomes and paternally derived nuclear alleles can produce directional postzygotic isolation (Burton & Barreto, 2012). Alternatively, species-specific maternal transcripts, proteins or cytoplasmic components deposited in the egg may interact poorly with heterospecific paternal chromosomes. All four interspecific crossing directions nevertheless produced at least some viable offspring, confirming incomplete reproductive isolation. The backcross-like ancestry observed in field-collected individuals further indicates that at least some hybrid descendants survive and reproduce, although the precise parental direction and pedigree of the natural crosses could not be inferred from the current nuclear markers. Mitochondrial haplotyping of field-collected hybrids would provide a direct test of whether introgression is biased toward crosses involving *B. mucronatus* maternal parents.

Natural introgression between *B. xylophilus* and *B. mucronatus* may have important evolutionary and disease-management consequences. These species share similar life histories and vector associations but differ markedly in pathogenicity, with *B. xylophilus* causing pine wilt disease and *B. mucronatus* generally exhibiting little or no virulence (Kanzaki & Futai, 2006; Mamiya & Enda, 1979). Hybridization followed by repeated backcrossing provides a potential route for adaptive introgression, through which alleles associated with pathogenicity in *B. xylophilus* could enter predominantly *B. mucronatus* genomic backgrounds, or locally adaptive alleles from *B. mucronatus* could be incorporated into *B. xylophilus* backgrounds (Rius & Darling, 2014).

Such gene flow could potentially alter traits relevant to invasion and disease emergence, including pathogenicity, thermal tolerance, host use, and vector association. Hybridization and introgression have contributed to the emergence of novel genotypes in other plant pathogen systems, in some cases combining pathogenicity-associated traits with broader environmental tolerances or host ranges (Stukenbrock, 2013). The strongly asymmetric ancestry observed in several field-collected hybrids in the present study indicates that repeated backcrossing provides a genomic pathway through which such trait combinations could arise. However, the present data do not identify the introgressed genomic regions or demonstrate that the detected hybrids possess enhanced virulence, fitness, or environmental tolerance. Adaptive introgression should be regarded as a biologically plausible and management-relevant consequence of the observed gene flow, rather than as an outcome established by this study.

These potential consequences may be particularly relevant in Europe, where *B. m. kolymensis* is widely distributed (Braasch, 2001; Magnusson & Kulinich, 1996). Our crossing experiments demonstrated incomplete reproductive isolation between *B. xylophilus* and *B. m. kolymensis*, although reproductive compatibility varied markedly between cross directions and among reproductive components. Crosses with *B. xylophilus* as the maternal parent produced relatively large numbers of eggs with reduced hatchability, whereas the reciprocal cross produced fewer eggs with high hatchability. Thus, the present results demonstrate that viable hybrid offspring can be generated in both directions, with variable compatibility. If *B. xylophilus* becomes established in regions where *B. m. kolymensis* occurs, hybridization followed by repeated backcrossing could provide a route for introgression between invasive and locally adapted genomes. Such introgression could potentially combine pathogenicity-associated alleles from *B. xylophilus* with traits present in European *B. m. kolymensis* populations, including adaptation to cooler environments, thereby generating genotypes with ecological properties not represented by either parental lineage. Comparable processes in other plant–pathogen systems have generated novel genetic combinations associated with changes in host range, environmental tolerance, or pathogenicity (Stukenbrock, 2013). However, we did not measure the pathogenicity, thermal performance or long-term fertility of *B. xylophilus* × *B. m. kolymensis* hybrids, and the evolutionary success of such genotypes in Europe remains uncertain. Nevertheless, our findings reinforce the importance of preventing the establishment of *B. xylophilus* in Europe and indicate that monitoring programs in potential contact zones should incorporate multilocus genomic surveillance, because advanced backcrosses may be indistinguishable from parental species based on morphology or single-locus markers. Experimental evaluation of hybrid pathogenicity, thermal tolerance, successive-generation fertility, and compatibility with European vector species will be essential for incorporating introgression risk into future invasion and disease distribution models.

This study had several limitations. First, sampling was restricted to a single locality over two consecutive years, limiting the geographic generality of the observed pattern. However, Shiojiri represents a recent secondary contact zone, where invasive *B. xylophilus* became established in an area previously occupied by native *B. mucronatus*. This setting may have been particularly suitable for detecting recent hybridization and backcrossing. Because field-collected individuals were selected primarily from beetle-derived populations in which both species had been detected, the nine hybrids identified among 342 genotyped nematodes should not be interpreted as an unbiased estimate of hybrid frequency within the wider population. Broader sampling across recent invasion fronts and long-established sympatric regions will be needed to assess geographic variation in hybridization. Second, although MIG-seq is a reduced-representation approach, it enabled a relatively large number of laboratory and field-collected individuals to be genotyped within a common framework. Empirically calibrated diagnostic SNPs distinguished major hybrid classes, while analyses based on 2,756 genome-wide SNPs provided complementary support. Thus, MIG-seq was well suited to detecting recent hybridization and broadly distinguishing balanced from parentally biased ancestry. However, it could not resolve introgressed tract lengths, exact backcross generations, or specific transferred genes. Accordingly, F1-like and advanced-backcross-like assignments should be interpreted as ancestry categories rather than exact pedigrees. The empirical HIest framework was also based on a limited set of laboratory strain combinations and would benefit from validation using additional parental strains. Finally, the maternal origin, geographic origin, fitness, pathogenicity, thermal tolerance, and vector associations of the natural hybrids remain unresolved and will require mitochondrial markers, denser genomic data, and experimental phenotyping. Future studies integrating denser genomic data, mitochondrial ancestry, and experimental phenotyping will be needed to determine the direction, persistence, and functional consequences of introgression.

In conclusion, by combining pedigree-calibrated ancestry inference with complementary genome-wide analyses, we demonstrated that hybridization between invasive *B. xylophilus* and native *B. mucronatus* is an ongoing and recurrent evolutionary process in natural populations in Japan, extending beyond F1 formation into advanced backcrossing. The incomplete reproductive isolation observed in our laboratory crosses provides a plausible mechanistic basis for this natural gene flow and establishes *Bursaphelenchus* as a tractable system for investigating hybridization and introgression during biological invasion. The failure of a single-locus marker to detect some advanced backcrosses shows that cryptic introgression can be overlooked by conventional surveillance. Such gene flow may generate novel combinations of pathogenicity, environmental tolerance, host use, or vector association, although these phenotypic consequences remain to be tested. As climate change and range expansion increase opportunities for secondary contact, the *Bursaphelenchus* system offers a valuable framework for understanding and managing the evolutionary risks of hybridization in economically important forest pests.

## Acknowledgments

We thank Mitsuteru Akiba and Takuya Aikawa (FFPRI) for sharing the strain *B. mucronatus*. This work was funded by grant from JSPS KAKENHI (No. 25K02027 to R.S.) and JST FOREST (No. JPMJFR210A to R.S.) The computation was performed using Research Center for Computational Science, Okazaki, Japan (Project: NIBB, 25-IMS-C330).

## Author Contribution

Yuzuki Ikeda and Ryoji Shinya conceived of the experiments.

Yuzuki Ikeda and Ryoji Shinya were involved in all aspects of the project, Naoko Ishikawa and Yoshihisa Suyama were contributed to sequence data analysis, and Kenichi Yanagisawa was contributed to fieldwork. Yuzuki Ikeda wrote the first draft of the manuscript. Naoko Ishikawa, Yoshihisa Suyama, Kenichi Yanagisawa, and Ryoji Shinya edited the manuscript.

## Notes

### Competing Interest Statement

The authors have declared no competing interest.

