## Supporting Information for "Genome-wide evidence for cryptic multi-generational introgression between invasive *Bursaphelenchus xylophilus* and native *B. mucronatus*"

**Supplemental Methods**

**Nematode culture and production of laboratory hybrids**

All nematode strains were maintained at 25°C on malt extract agar (MEA; Difco Laboratories, Detroit, MI, USA) containing 100 μg/mL chloramphenicol (Fujifilm Wako Pure Chemical, Osaka, Japan) and inoculated with the BC-3 strain of *Botrytis cinerea*.

Laboratory hybrids of known pedigree were obtained by isolating individual L4-stage females and adult males from mixed cultures using a fine needle under a stereomicroscope. To ensure virginity, isolated individuals were maintained for 2 days on 1/10-diluted MEA containing 4% agar and 100 μg/mL chloramphenicol and inoculated with *Saccharomyces cerevisiae* (hereafter, "1/10 MEA–yeast plates") in 60-mm Petri dishes, following Shinya et al. (2014). Single-pair matings were established by transferring one virgin female and one virgin adult male onto a fresh 1/10 MEA–yeast plate. After egg deposition began, females were transferred to fresh plates every 2 days to maintain pedigrees separately.

F1 offspring were individually isolated after hatching and reared to adulthood. First-generation backcrosses (B1) were generated by mating virgin F1 adults with virgin individuals from the parental taxa using the same procedure. Second-generation backcrosses (B2) were similarly generated by mating virgin B1 individuals with parental individuals. All laboratory-generated hybrids were subjected to MIG-seq analysis and used as pedigree-confirmed reference classes (Table S2).

**Nematode extraction and species-specific screening of beetle-derived populations**

Nematodes were extracted separately from individual *M. alternatus* adults using a modified Baermann funnel method (Maehara et al., 2020). Each beetle was homogenized for 10 s in 40 mL distilled water using a commercial blender (Waring, Stamford, CT, USA); the resulting suspension was transferred to a Baermann funnel lined with a single layer of tissue paper (Kimwipes; Kimberly-Clark, Irving, TX, USA). After overnight incubation at room temperature (approximately 20–25°C), nematodes that had migrated through the tissue were collected from the funnel stem and counted under a stereomicroscope (SMZ 1500; Nikon, Tokyo, Japan). When nematode densities were too high for direct counting, suspensions were serially diluted and subsamples were counted to estimate the total nematode load per beetle. The nematodes recovered from each beetle were treated as a single population for subsequent molecular analyses.

For initial species screening, approximately 100 nematodes were randomly selected from each beetle-derived population and pooled in a 0.2-mL PCR tube containing 50 μL DirectPCR Lysis Reagent (Viagen Biotech, Los Angeles, CA, USA). Samples were subjected to five freeze–thaw cycles using liquid nitrogen to facilitate disruption of nematode tissues. Genomic DNA was extracted by adding 10 μL Proteinase K solution (20 mg/mL; Nacalai Tesque, Kyoto, Japan), followed by incubation at 60°C for 2 h and enzyme inactivation at 95°C for 10 min. When tissues were not completely digested, an additional 5–10 μL Proteinase K solution was added and incubation at 60°C was continued for 1 h.

Species identification was performed using diagnostic primer sets targeting species-specific regions of *B. xylophilus* and *B. mucronatus* (Li et al., 2021; Matsunaga et al., 2019; Matsunaga & Togashi, 2004; Table S3). PCR reactions were performed in a total volume of 25 μL containing 0.5 μL genomic DNA, 0.125 μL AmpliTaq Gold DNA Polymerase (5 U/μL; Thermo Fisher Scientific, Waltham, MA, USA), 2.5 μL 10× PCR Buffer II, 0.5 μL dNTP mix,

1.5 μL $MgCl_2$ solution, 0.5 μL each of forward and reverse primers, and 18.875 μL nuclease-free water. Thermal cycling was performed using a ProFlex PCR System (Applied Biosystems) with an initial denaturation at 94°C for 5 min; 35 cycles of 94°C for 30 s, primer-specific annealing at 55.9°C or 48.0°C for 30 s, and 72°C for 1 min; followed by a final extension at 72°C for 6 min.

PCR products were separated on 1.5% agarose gels containing GreenView Nucleic Acid Gel Stain (Applied Biological Materials, Richmond, BC, Canada) in 1× TAE (Tris base, acetic acid, and ethylenediaminetetraacetic acid) buffer and visualized using blue-light transillumination. Fragment sizes were estimated using a 100-bp DNA Ladder (Takara Bio). Populations showing diagnostic amplification for both *B. xylophilus* and *B. mucronatus* were selected for individual-level MIG-seq analysis.

For these mixed-species populations, 96 individual nematodes were randomly isolated into separate PCR tubes containing 5 μL DirectPCR Lysis Reagent and 1 μL Proteinase K solution (20 mg/mL). DNA extraction followed the freeze–thaw and Proteinase K digestion procedure described above. Individuals with sufficient DNA quality and quantity were subsequently selected for MIG-seq analysis.

**MIG-seq library preparation and read preprocessing**

MIG-seq libraries were prepared following Suyama & Matsuki (2015) and Suyama et al. (2022). The first PCR was performed using MIG-seq primer set 1. The sequences of the forward and reverse ISSR primers are provided in Table S7.

Products of the first PCR were purified using a magnetic bead-based cleanup method following Hosomichi et al. (2013). A second PCR was performed to add sample-specific index sequences and Illumina flow cell adapter sequences to the ISSR amplicons generated in the first PCR.

Reactions for the second PCR were conducted in a total volume of 6 µL containing 1.6 µL purified first-PCR product, 0.12 µL PrimeSTAR GXL DNA Polymerase (1.25 U/µL; Takara Bio), 1.2 µL 5× PrimeSTAR GXL Buffer ($Mg^{2+}$ plus; Takara Bio), 0.48 µL dNTP mix (2.5 mM each; Takara Bio), 2.4 µL forward and reverse primer mixture (0.5 µM each), and 0.2 µL nuclease-free water. Thermal cycling was performed using a MiniAmp Plus Thermal Cycler (Applied Biosystems) for 12 cycles of 98°C for 10 s, 54°C for 15 s, and 68°C for 1 min.

The quality of the second-PCR products was assessed by microchip electrophoresis using a MultiNA system (Shimadzu, Kyoto, Japan) with a DNA-2500 reagent kit. Subsequently, 2 µL of the second-PCR product from each sample was pooled into a single tube, and the pooled products were purified by magnetic bead-based cleanup (Hosomichi et al., 2013).

The custom adapter sequences used for read trimming were as follows:

PrefixMIGAdapterF

CAGAGATCGGAAGAGCGTCGTGTAGGGAAAGA

PrefixMIGAdapterR

GTCAGATCGGAAGAGCACACGTCTGAACTCCAGTCAC

Raw sequence reads were quality filtered using Trimmomatic v0.39 (Bolger et al., 2014). Because sequencing data were generated on both MiSeq and DNBSEQ-G400 platforms, trimming parameters were optimized separately according to read length. MiSeq reads were processed using the parameters CROP = 77, HEADCROP = 6, and MINLEN = 51. DNBSEQ-G400 reads were processed using the parameters CROP = 100, HEADCROP = 17, and MINLEN = 51, together with the custom adapter file described above.

Paired and unpaired reads that passed quality filtering were retained. Unpaired reads generated during trimming were concatenated with their corresponding paired reads; and forward and reverse reads were subsequently combined into a single FASTQ file for each individual. Samples yielding fewer than 50,000 reads after trimming were excluded from subsequent analyses.

### Detailed procedures for ADMIXTURE, PCA, and NeighborNet analyses

#### *Supervised ADMIXTURE analysis*

For the *B. xylophilus*–*B. m. mucronatus* comparison, quality-filtered reads were assembled using ipyrad v0.9.102 using *B. xylophilus* as the reference genome (GCA_904066235.2_BXYJv5). The maximum heterozygosity per locus was set to 0.6, and loci were retained when present in at least 305 of the initial 436 individuals.

Diagnostic SNPs were selected using the same general criteria as those used for HIest analysis. Because sequencing depth and missing data rates varied among laboratory and field samples, SNPs differentiated between the parental taxa were further filtered across the combined laboratory and field dataset to retain loci that were sufficiently represented across samples. SNPs were retained according to the following criteria: $\leq$ 25% missing data; allele frequency divergence between parental taxa of $|\Delta| \geq 0.80$; at least one minor allele in the parental dataset; and a minimum physical separation of 1 kb between retained SNPs to reduce linkage. Individuals for which no informative loci remained after filtering were excluded.

The resulting dataset contained 51 diagnostic SNPs for 436 individuals. For graphical presentation, individuals with > 50% missing genotypes across the final 51-SNP panel were omitted, resulting in a displayed dataset of 411 individuals (122 laboratory samples, 120 field-collected individuals from 2023, and 169 field-collected individuals from 2024). In supervised

ADMIXTURE analysis (K = 2), field-collected individuals were analyzed without predefined ancestry assignments. Ancestry coefficients obtained from ADMIXTURE analysis were used as complementary indicators of genomic ancestry rather than as the primary criterion for assigning hybrid generations.

*PCA*

PCA was performed using a broader genome-wide SNP dataset independently filtered from the diagnostic SNP panel. SNP loci with > 30% missing genotypes were excluded, followed by removal of individuals with > 50% missing genotypes. The final dataset contained 2,756 SNPs from 416 individuals, comprising 122 laboratory individuals, 125 field-collected individuals from 2023, and 169 field-collected individuals from 2024.

PCA was performed using the glPca function in the R package *adegenet*. The proportion of variance explained by each principal component (PC) was calculated from the corresponding eigenvalues, and individual scores were visualized as two-dimensional PC1–PC2 plots using the R package ggplot2.

*NeighborNet analysis*

NeighborNet analysis was conducted using the same dataset of 2,756 SNPs and 416 individuals used for PCA. Pairwise genetic distances among individuals were calculated using uncorrected p-distances, and NeighborNet networks were constructed using SplitsTree v6.6.1. Ambiguous sites were handled using the MatchStates option.

**Detailed chemotaxis assay procedure**

Chemotaxis assays were performed in 90-mm Petri dishes containing 4% agar. Approximately 100 virgin males or females suspended in approximately 10 µL water were placed at the center of each plate within a 3-cm-diameter buffer zone.

Moments before the water containing the test nematodes was completely absorbed into the agar, two 1-µL drops of 1 M sodium azide were placed on opposite sides of the plate, approximately 1 cm from the plate edge. Two 10-µL drops of dd$H_2O$ were then placed on the inner surface of the Petri dish lid directly above the sodium azide positions. Ten virgin individuals of the opposite sex were transferred into one of the water drops to provide a volatile cue, while the second drop contained dd$H_2O$ alone (control). Sodium azide was used to immobilize test nematodes after reaching the cue or control zone, thereby minimizing subsequent redistribution and potential effects of behavioral adaptation.

Plates were placed in a small enclosed box to minimize the influence of light and incubated for 6 h. After incubation, immobilized test nematodes that had moved outside the central 3-cm buffer zone were scored according to whether they accumulated on the nematode-cue or control side.

Chemotactic preference was calculated as: CI = ([Number of nematodes in the test cue zone] – [Number of nematodes in the control zone])/([Number of nematodes in the test cue zone] + [Number of nematodes in the control zone]), where positive CI values indicate attraction toward the nematode-derived volatile cue and negative values indicate avoidance.

**Detailed fecundity and hatchability assay procedure**

For post-mating reproductive-isolation assays, one virgin female was placed together with three virgin males on a 1/10 MEA–yeast plate to increase the probability of successful mating and sperm transfer.

The following conspecific crosses were examined: *B. xylophilus* T4 female × T4 male, *B. m. mucronatus* Un-1 female × Un-1 male, and *B. m. kolymensis* TCS00 female × TCS00 male. Reciprocal interspecific crosses comprised *B. xylophilus* T4 female × *B. m. mucronatus* Un-1 male, *B. m. mucronatus* Un-1 female × *B. xylophilus* T4 male, *B. xylophilus* T4 female × *B. m. kolymensis* TCS00 male, and *B. m. kolymensis* TCS00 female × *B. xylophilus* T4 male.

Mated females were monitored daily and transferred to fresh 1/10 MEA–yeast plates every 24 h to separate eggs according to oviposition date. Daily transfers continued until females ceased laying eggs, typically within 5–10 days after mating. After each female had been removed, all eggs on the plate were counted under a stereomicroscope (SMZ 1500; Nikon). For each female, total fecundity was calculated as the cumulative number of eggs laid during the entire reproductive period.

To assess offspring viability, plates containing eggs were maintained at 25°C and hatching was scored at 24 h after oviposition. Hatchability was calculated as the number of hatched eggs divided by the total number of eggs laid. Trials in which the experimental female escaped during the assay were excluded because complete reproductive output could not be determined.

The tail morphology of the female F1 was observed using a differential interference contrast microscope (Bx53; Olympus).

### Supplemental Results

*Detailed classification performance of theoretical and empirical HIest frameworks*

Laboratory individuals of known pedigree occupied class-specific regions of S–H space in both species comparisons. Although the ordering of parental, F1, B1, and B2 classes was broadly consistent with theoretical expectations, empirical distributions shifted relative to the corresponding theoretical positions.

The most consistent deviation was a reduction in interspecific heterozygosity. This pattern was particularly pronounced in F1 individuals but was also evident in several backcross classes. Class-specific theoretical expectations, empirical means, and SDs are reported in Table 1.

Classification performance was evaluated using laboratory individuals of known pedigree that were independent of the parental reference individuals used for HIest estimation. In the *B. xylophilus–B. m. kolymensis* dataset, theoretical classification correctly assigned 48 of 76 individuals (63.2%), whereas empirical classification evaluated using LOOCV correctly assigned 59 of 76 individuals (77.6%). In the *B. xylophilus–B. m. mucronatus* dataset, theoretical classification correctly assigned 60 of 95 individuals (63.2%), whereas empirical LOOCV classification correctly assigned 77 of 95 individuals (81.1%). Overall and class-specific classification accuracies are provided in Table S8.

The improvement in overall classification accuracy under the empirical framework was driven primarily by improved classification of F1 and some backcross classes. However, improvement levels were not uniform across all classes, and classification accuracy remained low or decreased for some parental and backcross classes. These class-specific estimates should be interpreted with caution because the numbers of pedigree-confirmed individuals differed among classes and were very low for some classes. Overall, the use of empirical class centers and class-specific variation improved classification performance across the independent validation datasets under the observed marker and missing-data conditions.

*Species composition of beetle-derived nematode populations*

In 2023, 115 adult *M. alternatus* beetles were collected. Species-specific PCR detected both *B. xylophilus* and *B. mucronatus* in 31 beetle-derived populations (27.0%), *B. xylophilus* alone in 23 populations (20.0%), and *B. mucronatus* alone in 39 populations (33.9%). Neither species was detected in the remaining 22 beetles (19.1%).

In 2024, 126 beetles were collected. Both species were detected in 20 beetle-derived populations (15.9%), *B. xylophilus* alone in 48 populations (38.1%), and *B. mucronatus* alone in 34 populations (27.0%). Neither species was detected in the remaining 24 beetles (19.0%). Beetle-level collection information, nematode abundance, species-screening results, and recovery of hybrid individuals are provided in Table S6.

*Detailed HIest assignments of field-collected individuals*

Most field-collected individuals occupied regions of S–H space corresponding to one of the parental taxa, fewer occupied intermediate or backcross-like positions.

Nine individuals were classified as high-confidence hybrids because their HIest assignments were broadly concordant with the ancestry patterns obtained using ADMIXTURE and their genomic positions in NeighborNet analysis and PCA. These comprised 4-g, 4-i, and 14-h from 2023 and 3-f, 5-r, 5-t, 12-k, 15-f, and 15-i from 2024.

Additional individuals were assigned to, or located close to, hybrid reference classes by HIest alone. These individuals were retained as HIest-only candidates rather than included among the high-confidence hybrids. Individual S and H estimates, numbers of informative loci, empirical class assignments, standardized distances, complementary genomic results, and final status are provided in Table S9.

*Detailed supervised ADMIXTURE results*

Supervised ADMIXTURE analysis based on 51 diagnostic SNPs included 411 individuals: 122 laboratory individuals, 120 field-collected individuals from 2023, and 169 field-collected individuals from 2024.

Laboratory parental individuals showed ancestry coefficients close to the corresponding parental extremes, whereas pedigree-confirmed laboratory hybrids showed intermediate or parentally biased ancestry profiles broadly consistent with their pedigrees.

Among the 2023 field hybrids, individuals 4-g and 14-h showed relatively balanced *B. xylophilus* ancestry coefficients of approximately 0.54 and 0.56, respectively. Individual 4-i showed a strongly asymmetric ancestry profile that was consistent with advanced backcrossing.

Among the 2024 hybrids, individuals 5-t and 5-r showed similar, relatively balanced *B. xylophilus* ancestry coefficients of approximately 0.53 and 0.53, respectively. Individuals 3-f, 12-k, 15-f, and 15-i showed strongly asymmetric *B. xylophilus* ancestry coefficients of approximately 0.02, 0.07, 0.04, and 0.06, respectively, indicating advanced backcrossing toward *B. mucronatus*. Individual ancestry coefficients for all samples are provided in Table S9.

*Detailed NeighborNet and PCA results*

NeighborNet analysis based on 2,756 genome-wide SNPs separated the parental taxa into differentiated regions of the network. Pedigree-confirmed laboratory F1 hybrids occupied intermediate positions, whereas laboratory backcrosses were generally displaced toward the recurrent parental group.

Field-collected individuals with relatively balanced ancestry were also positioned between the parental taxa, while putative advanced backcrosses were located closer to one of the parental regions. Thus, the network broadly supported the distinction between F1-like and advanced-backcross-like field individuals inferred from HIest and ADMIXTURE analyses.

PCA based on the same dataset produced a generally consistent pattern (Figure S2). Parental samples formed differentiated clusters, laboratory-generated hybrids occupied intermediate or parentally displaced positions according to pedigree, and high-confidence field hybrids were distributed between or near the parental clusters. Individual PC1 and PC2 scores are provided in Table S9.

*Detailed chemotaxis responses*

Male responses to female cues did not differ significantly among conspecific and heterospecific combinations for either the *B. xylophilus–B. m. mucronatus* comparison ($F_{3, 15}$ = 1.182, $P$ = 0.3498) or the *B. xylophilus–B. m. kolymensis* comparison ($F_{3, 17}$ = 1.684, $P$ = 0.2082).

Mean chemotaxis indices for *B. xylophilus* males responding to *B. xylophilus*, *B. m. mucronatus*, and *B. m. kolymensis* female cues were 0.73 ± 0.09, 0.60 ± 0.08, and 0.40 ± 0.14, respectively (mean ± SE). No pairwise comparison was significant.

Female responses to male cues also did not differ among treatments for either the *B. xylophilus–B. m. mucronatus* comparison ($F_{3, 8}$ = 2.158, $P$ = 0.1711) or the *B. xylophilus–B. m. kolymensis* comparison ($F_{3, 8}$ = 0.7445, $P$ = 0.5551). Chemotaxis indices ranged from 0.25 to 0.96 across female response combinations. Crossing combinations and replication numbers are provided in Table S4.

*Detailed reproductive output of conspecific and interspecific crosses*

Eggs were produced in 10 of 12 *B. xylophilus* conspecific trials, 15 of 23 *B. m. mucronatus* conspecific trials, and 12 of 13 *B. m. kolymensis* conspecific trials. In interspecific crosses, eggs were produced in 20 of 23 *B. xylophilus* female × *B. m. mucronatus* male trials, 21 of 23 reciprocal trials, 10 of 11 *B. xylophilus* female × *B. m. kolymensis* male trials, and 10 of 15 reciprocal trials.

Among females that produced eggs, mean fecundity was 27.0 ± 4.04 eggs for *B. xylophilus*, 15.8 ± 2.32 for *B. m. mucronatus*, and 43.3 ± 7.38 for *B. m. kolymensis* conspecific crosses. Mean fecundity was 8.1 ± 0.91 eggs for *B. xylophilus* female × *B. m. mucronatus* male, 5.3 ± 1.12 for the reciprocal cross, 25.6 ± 4.99 for *B. xylophilus* female × *B. m. kolymensis* male, and 3.6 ± 0.85 for the reciprocal cross.

Mean hatchability was 97.5 ± 6.3%, 87.8 ± 11.5%, and 86.4 ± 16.5% in the three conspecific crosses, respectively. Among interspecific crosses, hatchability was 24.1 ± 26.6% for *B. xylophilus* female × *B. m. mucronatus* male and 76.3 ± 28.0% in the reciprocal direction. Hatchability was 56.0 ± 20.7% for *B. xylophilus* female × *B. m. kolymensis* male and 92.8 ± 12.0% in the reciprocal direction.

Complete descriptive statistics, model summaries, and pairwise contrasts are provided in Table S5.

**Supplemental figures**

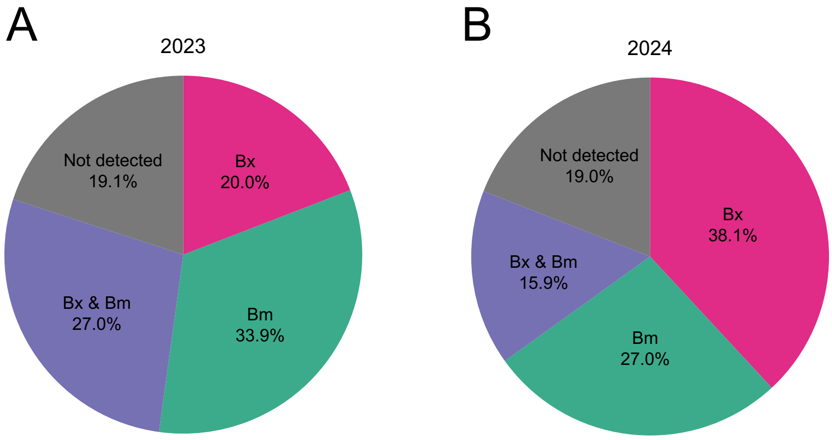


**Figure S1.** Species identification of nematodes carried by *Monochamus alternatus* collected in 2023 and 2024. Proportional compositions of nematode species detected from each of (A) 115 individual *M. alternatus* beetles collected in 2023 and (B) 126 individual *M. alternatus* beetles collected in 2024. Beetle-level collection information, nematode abundance, and species-screening results are provided in Table S6. Bx, *Bursaphelenchus xylophilus*; Bm, *B. mucronatus*.

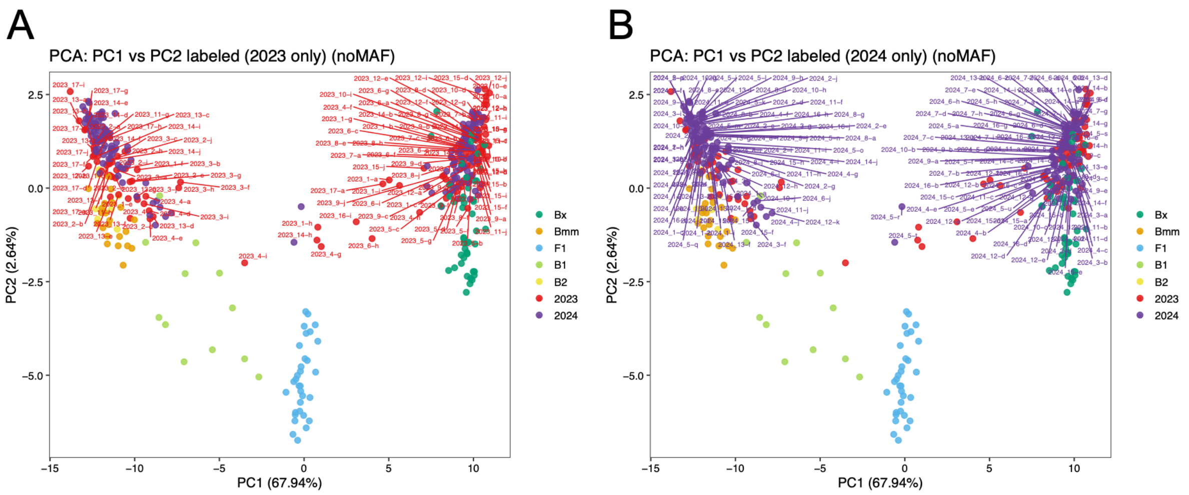


**Figure S2.** Principal component analysis plots of laboratory and field samples based on 2,756 independently filtered genome-wide single-nucleotide polymorphisms (SNPs). Colors indicate genetic classes represented by laboratory parental and known-pedigree hybrid individuals and field-collected individuals sampled in (A) 2023 and (B) 2024. Only field samples are labeled in each panel. Principal component 1 (PC1), which explained 67.94% of the variation, primarily reflected genetic differentiation between Bx and Bmm, whereas PC2, which explained 2.64%, captured additional variation among ancestry classes.

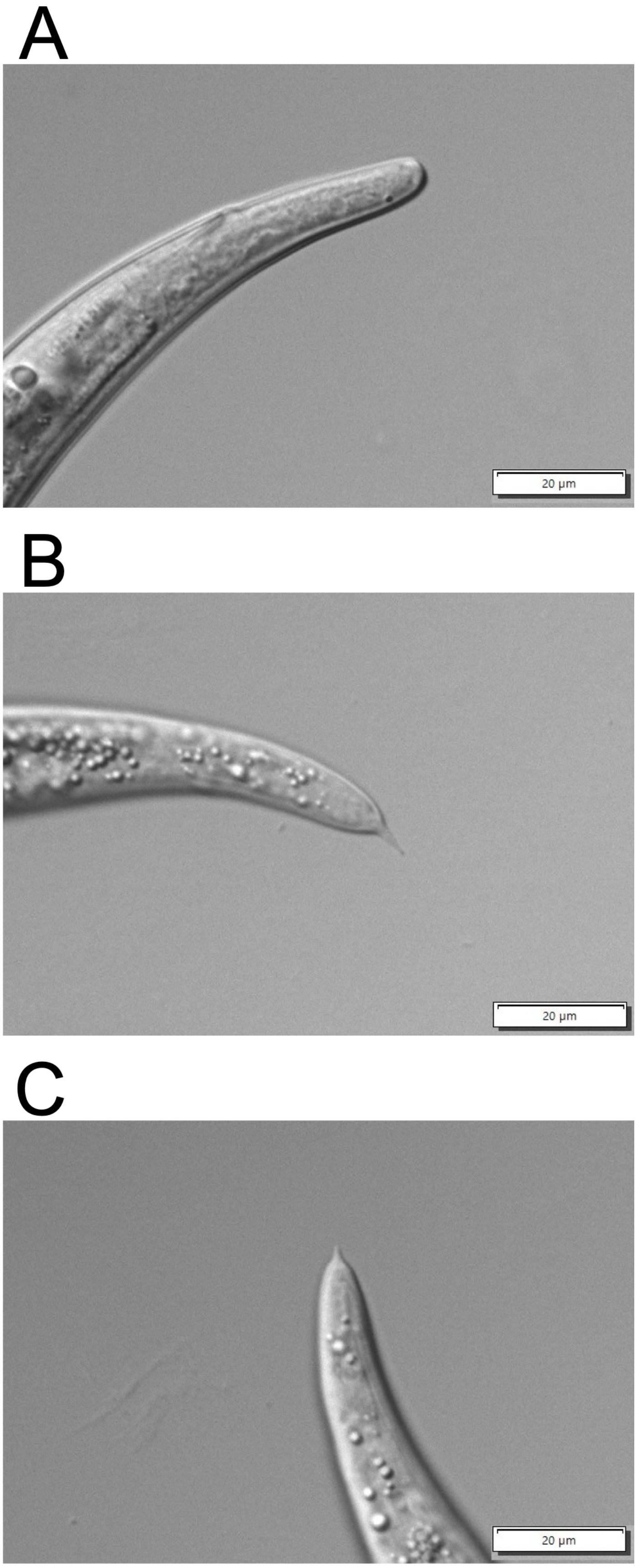


357

**Figure S3.** Tail morphology in purebreds and hybrids. Female tail morphology used for morphological species identification of (A) *B. xylophilus* (T4, round tail) and (B) *B. mucronatus* (Un-1, mucronate tail). (C) Tail of laboratory-generated F1 between an Un-1 female and a T4 male.

**Supplemental tables**

**Table S1.** *Bursaphelenchus xylophilus* (Bx), *Bursaphelenchus mucronatus mucronatus* (Bmm), and *Bursaphelenchus mucronatus kolymensis* (Bmk) strains used in genomic analyses and reproductive isolation experiments.

| Species | Sub-species | Isolate | Source | Location | Reference |
|---|---|---|---|---|---|
| Bx | | T4 | *Monochamus alternatus* | Iwate Pref. | Taga et al. (2011) |
| | | TBx#1 | *Pinus densiflora* | Hiroshima Pref. | Taga et al. (2011) |
| | | Ka4 | *Pinus densiflora* | Ibaraki Pref. | Shinya et al. (2021) |
| | | S10 | *Pinus densiflora* | Shimane Pref. | Shinya et al. (2012) |
| | | OKD-1 | *Pinus densiflora* | Shimane Pref. | Zhou et al. (2007) |
| | | C14-5 | *Monochamus alternatus* | Chiba Pref. | Fukuda et al. (1992) |
| | | NG2 | *Monochamus alternatus* | Nagano Pref. | |
| | | S6-1 | *Monochamus alternatus* | Ibaraki Pref. | Takemoto and Futai (2007) |
| | | TU3-2 | *Pinus densiflora* | Tokyo Pref. | |
| | | TN1 | *Pinus amamianal* | Kagoshima Pref. | |
| Bm | Bmm | Un-1 | *Pinus densiflora* | Nagasaki Pref. | Taga et al. (2011) |
| | | NG-1 | *Pinus densiflora* | Nagano Pref. | Mastunaga et al. (2019) |
| | | Ioujima | *Pinus thunbergii* | Kagoshima Pref. | Mastunaga et al. (2019) |
| | Bmk | Kitazawa 2 | *Monochamus nitens* | Yamanashi Pref. | |
| | | H30 | *Monochamus rosenmuelleri* | Hokkaido Pref. | |
| | | H110 | *Monochamus rosenmuelleri* | Hokkaido Pref. | |
| | | Srf | *Monochamus urussovi* | Hokkaido Pref. | Togashi et al. (2008) |
| | | TCS00 | *Pinus parviflora* | Fukushima Pref. | Taga et al. (2011) |
| | | TBm119 | *Monochamus saltuarius* | Hiroshima Pref. | Taga et al. (2011) |

**Table S2.** Laboratory-generated crosses among *B. xylophilus*, *B. m. mucronatus*, and *B. m. kolymensis* strains. Laboratory parental, F1, B1, and B2 individuals used to calibrate and validate the HIest-based ancestry-classification framework are shown. Different lowercase letters indicate crosses derived from different parental (P0) individuals.

| | | | | | | | | |
|---|---|---|---|---|---|---|---|---|
| **F1** | (TCS00-T4)a | (TCS00-T4)a | (TCS00-T4)a | (TCS00-T4)a | (TCS00-T4)a | (TCS00-T4)a | (TCS00-T4)b | (TCS00-T4)b |
| | (TCS00-T4)b | (TCS00-T4)b | (TCS00-T4)b | (TCS00-T4)b | (TCS00-T4)b | (TCS00-T4)b | (TCS00-T4)b | (TCS00-T4)c |
| | (TB#-1-Srf)d | (TB#-1-Srf)d | (TB#-1-Srf)d | (TB#-1-Srf)d | (TB#-1-Srf)d | (TB#-1-Srf)e | (TB#-1-Srf)e | (TB#-1-Srf)e |
| | (Ka4-loujima)f | (Ka4-loujima)f | (Ka4-loujima)f | (Ka4-loujima)f | (Ka4-loujima)f | (Ka4-loujima)f | (T4-loujima)g | (T4-loujima)g |
| | (T4-loujima)g | (T4-loujima)g | (S10-NG1)h | (S10-NG1)h | (S10-NG1)h | (S10-NG1)h | (S10-NG1)h | (S10-NG1)i |
| | (S10-NG1)j | (S10-NG1)j | (S10-NG1)j | (S10-NG1)j | (S10-NG1)j | (TB#-1-NG1)k | (TB#-1-NG1)k | (TB#-1-NG1)k |
| | (TB#-1-NG1)k | (TB#-1-NG1)k | (TB#-1-NG1)k | (TB#-1-NG1)l | (TB#-1-NG1)l | (TB#-1-NG1)l | (TB#-1-NG1)l | (TB#-1-NG1)l |
| **B1** | ((TCS00-T4)a-TCS00)a | ((TCS00-T4)a-TCS00)a | ((TCS00-T4)a-TCS00)a | ((TCS00-T4)a-TCS00)a | ((TCS00-T4)a-TCS00)a | ((TCS00-T4)a-TCS00)a | ((TCS00-T4)a-TCS00)a | ((TCS00-T4)a-TCS00)a |
| | ((TCS00-T4)b-TCS00)b | ((TCS00-T4)b-TCS00)b | ((TCS00-T4)b-TCS00)b | ((TCS00-T4)b-TCS00)b | ((TCS00-T4)b-TCS00)b | ((TCS00-T4)b-TCS00)b | ((TCS00-T4)b-TCS00)b | ((TCS00-T4)a-T4)c |
| | ((TCS00-T4)a-T4)c | ((TCS00-T4)a-T4)c | ((TCS00-T4)a-T4)c | ((TCS00-T4)a-T4)c | (loujima-(Ka4-loujima)f)d | (loujima-(Ka4-loujima)f)d | (loujima-(Ka4-loujima)f)d | (loujima-(Ka4-loujima)f)d |
| | ((TBx#1-Srf)e-Srf)e | ((TBx#1-Srf)e-Srf)e | ((TBx#1-Srf)e-Srf)e | ((TBx#1-Srf)e-Srf)e | ((TBx#1-Srf)e-Srf)e | ((TBx#1-Srf)e-Srf)e | ((TBx#1-Srf)e-Srf)e | ((TBx#1-Srf)e-Srf)e |
| | (NG1-(TBx#1-NG1)l)f | (NG1-(TBx#1-NG1)l)f | (NG1-(TBx#1-NG1)l)f | (NG1-(TBx#1-NG1)l)f | (NG1-(TBx#1-NG1)l)f | (NG1-(TBx#1-NG1)l)f | (NG1-(TBx#1-NG1)l)f | (NG1-(TBx#1-NG1)l)f |
| **B2** | (((TCS00-T4)a-TCS00)a-TCS00)a | (((TCS00-T4)a-TCS00)a-TCS00)a | (((TCS00-T4)a-TCS00)a-TCS00)a | (((TCS00-T4)a-TCS00)a-TCS00)a | (((TCS00-T4)a-TCS00)a-TCS00)a | (((TCS00-T4)a-TCS00)a-TCS00)a | (((TCS00-T4)a-TCS00)a-TCS00)a | (((TCS00-T4)a-TCS00)a-TCS00)a |
| | (((TCS00-T4)a-TCS00)a)-TCS00)b | (((TCS00-T4)b-TCS00)b-TCS00)c | (((TCS00-T4)b-TCS00)b-TCS00)c | (((TCS00-T4)b-TCS00)b-TCS00)c | (((TCS00-T4)b-TCS00)b-TCS00)c | (((TCS00-T4)b-TCS00)b-TCS00)c | (((TCS00-T4)b-TCS00)b-TCS00)c | (((TCS00-T4)b-TCS00)b-TCS00)c |
| | (((TBx#1-Srf)e-Srf)e-Srf)d | (((TBx#1-Srf)e-Srf)e-Srf)d | (((TBx#1-Srf)e-Srf)e-Srf)d | (((TBx#1-Srf)e-Srf)e-Srf)d | (((TBx#1-Srf)e-Srf)e-Srf)d | (((TBx#1-Srf)e-Srf)e-Srf)d | (((TBx#1-Srf)e-Srf)e-Srf)d | (((TBx#1-Srf)e-Srf)e-Srf)d |
| | ((NG1-(TBx#1-NG1)l)f-NG1)e | ((NG1-(TBx#1-NG1)l)f-NG1)e | ((NG1-(TBx#1-NG1)l)f-NG1)e | ((NG1-(TBx#1-NG1)l)f-NG1)e | ((NG1-(TBx#1-NG1)l)f-NG1)e | ((NG1-(TBx#1-NG1)l)f-NG1)e | ((NG1-(TBx#1-NG1)l)f-NG1)e | ((NG1-(TBx#1-NG1)l)f-NG1)e |

**Table S3.** Single-locus markers used for nematode species identification and hybrid detection. (A) Primers used for species identification in 2023; it is unclear whether these markers can discriminate hybrids (Matsunaga et al., 2019; Matsunaga & Togashi, 2004). (B) Hybrid-diagnostic primers used in 2024 (Li et al., 2021).

A

| Species | Origin | Sequence | Length of PCR product (bp) |
|---|---|---|---|
| **Bx** | mtCO1 | F:GATGAACTGTTTATCCTCCA<br>R:ACAACCAATTAAACCAATTC | 479 |
| **Bm** | ITS2<br>ITS2 to 28S rRNA gene | F:TCCGGCCATATCTCTACGAC<br>R:GTTTCAACCAATTCCGAACC | 210 |

B

| Species | Origin | Sequence | Length of PCR product (bp) |
|---|---|---|---|
| **Bx** | ITS1<br>5.8S | F:GATGATGCGATTGGTGACT<br>R:CAATTCACTGCGTTCTTC | 329 |
| **Bm** | ITS1<br>5.8S | F: TGCGCTGCGTTGAGTCGA<br>R: CAATTCACTGCGTTCTTC | 206 |

**Table S4.** Combinations and replication numbers for chemotaxis assays. Combinations of responding nematodes and opposite-sex chemical cues, together with the numbers of independent replicates, are shown for assays involving *B. xylophilus* strain T4, *B. m. mucronatus* strain Un-1, and *B. m. kolymensis* strain TCS00). Assays measured both male responses to female-derived cues and female responses to male-derived cues.

| Attraction of males to females | | Attraction of females to males | |
|---|---|---|---|
| Combinations (female - male) | Replication number | Combinations (female - male) | Replication number |
| Bx - Bx | 7 | Bx - Bx | 3 |
| Bmm - Bmm | 3 | Bmm - Bmm | 3 |
| Bmk - Bmk | 5 | Bmk - Bmk | 3 |
| Bx - Bmm | 5 | Bx - Bmm | 3 |
| Bmm - Bx | 4 | Bmm - Bx | 3 |
| Bx - Bmk | 6 | Bx - Bmk | 3 |
| Bmk - Bx | 3 | Bmk - Bx | 3 |

**Table S5.** Summary of reproductive performance in conspecific and interspecific crosses between *B. xylophilus* strain T4 and *B. m. mucronatus* strain Un-1 (A), and between *B. xylophilus* strain T4 and *B. m. kolymensis* strain TCS00 (B). Fecundity was summarized among egg-producing females, whereas hatchability was based on hatched and unhatched egg counts. Different letters indicate significant pairwise differences after adjustment for multiple comparisons.

A

| Combinations (female - male) | Trial number | Trial number which one or more eggs were laid | Number of eggs laid (mean ± SE) | | | | Hatchability rate (%) (mean ± SD) | | | |
|---|---|---|---|---|---|---|---|---|---|---|
| Bx - Bx | 12 | 10 | 27.0 | ± | 4.04 | a | 97.5 | ± | 6.3 | a |
| Bmm - Bmm | 23 | 15 | 15.8 | ± | 2.32 | b | 87.8 | ± | 11.5 | a |
| Bx - Bmm | 23 | 20 | 8.1 | ± | 0.91 | c | 24.1 | ± | 26.6 | b |
| Bmm - Bx | 23 | 21 | 5.3 | ± | 1.12 | c | 76.3 | ± | 28.0 | a |

B

| Combinations (female - male) | Trial number | Trial number which one or more eggs were laid | Number of eggs laid (mean ± SE) | | | | Hatchability rate (%) (mean ± SD) | | | |
|---|---|---|---|---|---|---|---|---|---|---|
| Bx - Bx | 12 | 10 | 27.0 | ± | 4.04 | a | 97.5 | ± | 6.3 | a |
| Bmk - Bmk | 13 | 12 | 43.3 | ± | 7.38 | a | 86.4 | ± | 16.5 | a |
| Bx - Bmk | 11 | 10 | 25.6 | ± | 4.99 | a | 56.0 | ± | 20.7 | b |
| Bmk - Bx | 15 | 10 | 3.6 | ± | 0.85 | b | 92.8 | ± | 12.0 | a |

**Table S6.** Numbers and species composition of nematodes isolated from individual *Monochamus alternatus* beetles in 2023 and 2024. Bx, *B. xylophilus*; Bm, *B. mucronatus*. Rows marked with a cross indicate that neither Bx nor Bm was detected by the initial species-screening assay.

| 2023 | | | | |
|---|---|---|---|---|
| Beetle number | Capture date | Log storage location | Nunber of nematodes | Nematode species detected by initial species screening |
| 1 | 6/10 | Kanagawa Pref. | 3400 | Bx |
| 2 | 6/15 | Kanagawa Pref. | 9400 | Bx & Bm |
| 3 | 6/15 | Kanagawa Pref. | 1000 | Bx |
| 4 | 6/16 | Kanagawa Pref. | 2100 | Bx |
| 5 | 6/16 | Kanagawa Pref. | 13100 | Bx |
| 6 | 6/16 | Kanagawa Pref. | 3300 | Bx & Bm |
| 7 | 6/18 | Kanagawa Pref. | 2000 | Bx & Bm |
| 8 | 6/19 | Kanagawa Pref. | 5400 | Bx |
| 9 | 6/21 | Kanagawa Pref. | 1100 | Bm |
| 10 | 6/22 | Kanagawa Pref. | 2733 | Bm |
| 11 | 6/22 | Kanagawa Pref. | 800 | Bm |
| 12 | 6/23 | Kanagawa Pref. | 7067 | Bm |
| 13 | 6/23 | Kanagawa Pref. | 667 | Bx |
| 14 | 6/23 | Kanagawa Pref. | 73 | × |
| 15 | 6/24 | Kanagawa Pref. | 3067 | Bx & Bm |
| 16 | 6/24 | Kanagawa Pref. | 533 | Bm |
| 17 | 6/24 | Kanagawa Pref. | 267 | Bx |
| 18 | 6/24 | Kanagawa Pref. | 1200 | Bx & Bm |
| 19 | 6/25 | Kanagawa Pref. | 5 | Bm |
| 20 | 6/25 | Kanagawa Pref. | 3133 | Bx |
| 21 | 6/26 | Kanagawa Pref. | 2 | Bm |
| 22 | 6/26 | Kanagawa Pref. | 267 | × |
| 23 | 6/26 | Kanagawa Pref. | 1400 | Bx & Bm |
| 24 | 6/26 | Kanagawa Pref. | 533 | Bm |
| 25 | 6/26 | Kanagawa Pref. | 10 | × |
| 26 | 6/26 | Kanagawa Pref. | 3 | × |
| 27 | 6/26 | Kanagawa Pref. | 2 | × |
| 28 | 6/26 | Kanagawa Pref. | 200 | Bm |
| 29 | 6/26 | Kanagawa Pref. | 8200 | Bm |
| 30 | 6/26 | Kanagawa Pref. | 12733 | Bm |
| 31 | 6/26 | Kanagawa Pref. | 15 | × |
| 32 | 6/26 | Kanagawa Pref. | 1067 | × |
| 33 | 6/26 | Kanagawa Pref. | 6933 | Bx |
| 34 | 6/27 | Kanagawa Pref. | 200 | Bm |
| 35 | 6/27 | Kanagawa Pref. | 5533 | Bx & Bm |
| 36 | 6/28 | Kanagawa Pref. | 333 | Bx & Bm |
| 37 | 6/28 | Kanagawa Pref. | 3467 | Bm |
| 38 | 6/28 | Kanagawa Pref. | 5 | Bm |
| 39 | 6/29 | Kanagawa Pref. | 3000 | Bm |
| 40 | 6/29 | Kanagawa Pref. | 3933 | Bm |
| 41 | 6/29 | Kanagawa Pref. | 2267 | Bm |
| 42 | 6/30 | Kanagawa Pref. | 10 | Bm |
| 43 | 6/30 | Kanagawa Pref. | 1400 | Bm |
| 44 | 6/30 | Kanagawa Pref. | 3067 | Bx & Bm |
| 45 | 7/3 | Kanagawa Pref. | 2733 | Bx & Bm |
| 46 | 7/5 | Kanagawa Pref. | 10200 | Bx & Bm |
| 47 | 7/5 | Kanagawa Pref. | 15533 | Bx & Bm |
| 48 | 7/5 | Kanagawa Pref. | 36 | Bm |
| 49 | 7/5 | Kanagawa Pref. | 400 | Bm |
| 50 | 7/18 | Nagano Pref. | 1000 | Bx |
| 51 | 7/18 | Nagano Pref. | 67 | Bx |
| 52 | 7/18 | Nagano Pref. | 2867 | Bx & Bm |
| 53 | 7/18 | Nagano Pref. | 533 | Bm |
| 54 | 7/18 | Nagano Pref. | 200 | × |
| 55 | 7/18 | Nagano Pref. | 6067 | Bx |
| 56 | 7/18 | Nagano Pref. | 400 | Bx & Bm |
| 57 | 7/18 | Nagano Pref. | 67 | Bm |
| 58 | 7/18 | Nagano Pref. | 933 | Bx & Bm |
| 59 | 7/18 | Nagano Pref. | 7 | Bx |
| 60 | 7/18 | Nagano Pref. | 67 | Bm |
| 61 | 7/18 | Nagano Pref. | 110 | Bm |
| 62 | 7/18 | Nagano Pref. | 4 | Bm |
| 63 | 7/18 | Nagano Pref. | 2000 | Bx & Bm |
| 64 | 7/18 | Nagano Pref. | 1000 | Bx & Bm |
| 65 | 7/18 | Nagano Pref. | 10 | Bx & Bm |
| 66 | 7/18 | Nagano Pref. | 3 | Bx & Bm |
| 67 | 7/18 | Nagano Pref. | 5 | Bm |
| 68 | 7/18 | Nagano Pref. | 2067 | Bm |
| 69 | 7/18 | Nagano Pref. | 1267 | Bx & Bm |
| 70 | 7/18 | Nagano Pref. | 533 | Bx |
| 71 | 7/18 | Nagano Pref. | 38933 | Bx & Bm |
| 72 | 7/18 | Nagano Pref. | 16067 | Bx |
| 73 | 7/18 | Nagano Pref. | 13400 | Bx & Bm |
| 74 | 7/18 | Nagano Pref. | 133 | Bx & Bm |
| 75 | 7/18 | Nagano Pref. | 25133 | Bx & Bm |
| 76 | 7/18 | Nagano Pref. | 400 | Bx & Bm |
| 77 | 7/18 | Nagano Pref. | 800 | Bx & Bm |
| 78 | 7/18 | Nagano Pref. | 200 | Bm |
| 79 | 7/18 | Nagano Pref. | 1200 | Bx & Bm |
| 80 | 7/31 | Nagano Pref. | 2 | Bm |
| 81 | 7/31 | Nagano Pref. | 30 | Bm |
| 82 | 7/31 | Nagano Pref. | 1 | × |
| 83 | 7/31 | Nagano Pref. | 0 | |
| 84 | 7/31 | Nagano Pref. | 0 | |
| 85 | 7/31 | Nagano Pref. | 0 | |
| 86 | 7/31 | Nagano Pref. | 1 | × |
| 87 | 7/31 | Nagano Pref. | 3 | × |
| 88 | 7/31 | Nagano Pref. | 200 | Bm |
| 89 | 7/31 | Nagano Pref. | 1 | Bm |
| 90 | 7/31 | Nagano Pref. | 0 | |
| 91 | 7/31 | Nagano Pref. | 14 | Bx |
| 92 | 7/31 | Nagano Pref. | 0 | |
| 93 | 7/31 | Nagano Pref. | 267 | Bm |
| 94 | 7/31 | Nagano Pref. | 12 | Bx |
| 95 | 7/31 | Nagano Pref. | 467 | Bm |
| 96 | 7/31 | Nagano Pref. | 1 | × |
| 97 | 7/31 | Nagano Pref. | 2533 | Bx & Bm |
| 98 | 8/1 | Nagano Pref. | 8 | × |
| 99 | 8/1 | Nagano Pref. | 0 | |
| 100 | 8/1 | Nagano Pref. | 123 | Bx & Bm |
| 101 | 8/1 | Nagano Pref. | 329 | Bx |
| 102 | 8/1 | Nagano Pref. | 205 | Bx |
| 103 | 8/1 | Nagano Pref. | 67 | Bx |
| 104 | 8/1 | Nagano Pref. | 200 | × |
| 105 | 8/2 | Nagano Pref. | 2 | Bm |
| 106 | 8/2 | Nagano Pref. | 1735 | Bx |
| 107 | 8/2 | Nagano Pref. | 3 | Bx |
| 108 | 8/2 | Nagano Pref. | 67 | Bm |
| 109 | 8/3 | Nagano Pref. | 1 | × |
| 110 | 8/3 | Nagano Pref. | 933 | Bx & Bm |
| 111 | 8/3 | Nagano Pref. | 1 | Bm |
| 112 | 8/4 | Nagano Pref. | 267 | Bx & Bm |
| 113 | 8/4 | Nagano Pref. | 667 | Bx |
| 114 | 8/9 | Nagano Pref. | 2 | × |
| 115 | 8/12 | Nagano Pref. | 467 | Bm |

| 2024 | | | | |
|---|---|---|---|---|
| Beetle number | Capture date | Log storage location | Nunber of nematodes | Nematode species detected by initial species screening |
| 1 | 6/10 | Kanagawa Pref. | 1760 | Bm |
| 2 | 6/10 | Kanagawa Pref. | 3540 | Bm |
| 3 | 6/10 | Kanagawa Pref. | 13600 | Bm |
| 4 | 6/10 | Kanagawa Pref. | 40 | Bm |
| 5 | 6/11 | Kanagawa Pref. | 160 | Bm |
| 6 | 6/11 | Kanagawa Pref. | 1820 | Bm |
| 7 | 6/12 | Kanagawa Pref. | 6780 | Bm |
| 8 | 6/12 | Kanagawa Pref. | 20 | × |
| 9 | 6/13 | Kanagawa Pref. | 640 | Bm |
| 10 | 6/14 | Kanagawa Pref. | 1080 | Bm |
| 11 | 6/14 | Kanagawa Pref. | 11580 | Bm |
| 12 | 6/14 | Kanagawa Pref. | 1840 | Bm |
| 13 | 6/15 | Kanagawa Pref. | 2680 | Bm |
| 14 | 6/15 | Kanagawa Pref. | 1260 | Bm |
| 15 | 6/15 | Kanagawa Pref. | 30 | Bm |
| 16 | 6/16 | Kanagawa Pref. | 1300 | Bm |
| 17 | 6/16 | Kanagawa Pref. | 260 | Bm |
| 18 | 6/16 | Kanagawa Pref. | 2020 | Bm |
| 19 | 6/16 | Kanagawa Pref. | 20 | Bm |
| 20 | 6/17 | Kanagawa Pref. | 60 | Bm |
| 21 | 6/17 | Kanagawa Pref. | 13820 | Bm |
| 22 | 6/17 | Kanagawa Pref. | 2100 | Bm |
| 23 | 6/19 | Kanagawa Pref. | 40 | × |
| 24 | 6/19 | Kanagawa Pref. | 9960 | Bm |
| 25 | 6/19 | Kanagawa Pref. | 400 | × |
| 26 | 6/20 | Kanagawa Pref. | 6240 | × |
| 27 | 6/20 | Kanagawa Pref. | 8600 | Bm |
| 28 | 6/20 | Kanagawa Pref. | 30400 | Bm |
| 29 | 6/20 | Kanagawa Pref. | 6200 | Bm |
| 30 | 6/20 | Kanagawa Pref. | 300 | Bm |
| 31 | 6/20 | Kanagawa Pref. | 500 | Bm |
| 32 | 6/20 | Kanagawa Pref. | 800 | Bm |
| 33 | 6/20 | Kanagawa Pref. | 16800 | Bx |
| 34 | 6/21 | Kanagawa Pref. | 3600 | Bx |
| 35 | 6/21 | Kanagawa Pref. | 1000 | Bm |
| 36 | 6/21 | Kanagawa Pref. | 200 | × |
| 37 | 6/21 | Kanagawa Pref. | 11600 | Bm |
| 38 | 6/21 | Kanagawa Pref. | 8800 | × |
| 39 | 6/22 | Kanagawa Pref. | 600 | × |
| 40 | 6/22 | Kanagawa Pref. | 23100 | Bx |
| 41 | 6/23 | Kanagawa Pref. | 300 | × |
| 42 | 6/23 | Kanagawa Pref. | 100 | × |
| 43 | 6/24 | Kanagawa Pref. | 0 | |
| 44 | 6/24 | Kanagawa Pref. | 15400 | Bx |
| 45 | 6/24 | Kanagawa Pref. | 2500 | Bx & Bm |
| 46 | 6/24 | Kanagawa Pref. | 3 | Bx |
| 47 | 6/25 | Kanagawa Pref. | 3000 | Bm |
| 48 | 6/25 | Kanagawa Pref. | 1600 | Bm |
| 49 | 6/25 | Kanagawa Pref. | 22000 | Bx |
| 50 | 6/25 | Kanagawa Pref. | 3800 | Bx |
| 51 | 6/25 | Kanagawa Pref. | 11900 | Bx & Bm |
| 52 | 6/27 | Kanagawa Pref. | 4900 | Bx |
| 53 | 6/29 | Kanagawa Pref. | 38 | × |
| 54 | 6/29 | Kanagawa Pref. | 500 | Bx & Bm |
| 55 | 6/29 | Kanagawa Pref. | 1400 | Bx & Bm |
| 56 | 6/30 | Nagano Pref. | 29800 | Bx |
| 57 | 6/30 | Nagano Pref. | 3500 | Bx |
| 58 | 7/1 | Nagano Pref. | 15200 | Bx |
| 59 | 7/2 | Nagano Pref. | 17 | Bx |
| 60 | 7/2 | Nagano Pref. | 11100 | Bx |
| 61 | 7/2 | Nagano Pref. | 500 | Bx |
| 62 | 7/3 | Kanagawa Pref. | 0 | |
| 63 | 7/3 | Kanagawa Pref. | 2000 | Bx |
| 64 | 7/4 | Nagano Pref. | 1 | Bx |
| 65 | 7/4 | Nagano Pref. | 8 | × |
| 66 | 7/4 | Nagano Pref. | 39100 | Bx & Bm |
| 67 | 7/4 | Nagano Pref. | 700 | Bx & Bm |
| 68 | 7/4 | Nagano Pref. | 78700 | Bx & Bm |
| 69 | 7/4 | Nagano Pref. | 44 | Bx & Bm |
| 70 | 7/5 | Kanagawa Pref. | 700 | Bx |
| 71 | 7/5 | Kanagawa Pref. | 3300 | × |
| 72 | 7/5 | Nagano Pref. | 53 | Bx |
| 73 | 7/6 | Kanagawa Pref. | 3400 | Bx |
| 74 | 7/7 | Kanagawa Pref. | 4 | Bx |
| 75 | 7/7 | Kanagawa Pref. | 14900 | Bx |
| 76 | 7/7 | Nagano Pref. | 2 | Bx |
| 77 | 7/7 | Nagano Pref. | 0 | |
| 78 | 7/8 | Nagano Pref. | 135 | Bx |
| 79 | 7/9 | Nagano Pref. | 0 | |
| 80 | 7/10 | Nagano Pref. | 0 | |
| 81 | 7/10 | Nagano Pref. | 0 | |
| 82 | 7/11 | Kanagawa Pref. | 13600 | Bx |
| 83 | 7/19 | Nagano Pref. | 300 | Bx |
| 84 | 7/19 | Nagano Pref. | 0 | |
| 85 | 7/21 | Nagano Pref. | 800 | Bx |
| 86 | 7/23 | Nagano Pref. | 8400 | Bx |
| 87 | 7/23 | Nagano Pref. | 400 | Bx |
| 88 | 7/23 | Nagano Pref. | 700 | Bx |
| 89 | 7/23 | Nagano Pref. | 600 | Bx |
| 90 | 7/24 | Nagano Pref. | 1 | Bx |
| 91 | 7/24 | Nagano Pref. | 700 | Bx |
| 92 | 7/24 | Nagano Pref. | 24900 | Bm |
| 93 | 7/24 | Nagano Pref. | 900 | Bx & Bm |
| 94 | 7/24 | Nagano Pref. | 3 | Bm |
| 95 | 7/24 | Nagano Pref. | 3300 | Bx & Bm |
| 96 | 7/25 | Nagano Pref. | 16 | Bx & Bm |
| 97 | 7/25 | Nagano Pref. | 35500 | Bx |
| 98 | 7/25 | Nagano Pref. | 2400 | Bx |
| 99 | 7/26 | Nagano Pref. | 1100 | Bx & Bm |
| 100 | 7/26 | Nagano Pref. | 200 | Bx |
| 101 | 7/26 | Nagano Pref. | 0 | |
| 102 | 7/26 | Nagano Pref. | 2300 | Bx |
| 103 | 7/28 | Nagano Pref. | 0 | |
| 104 | 7/28 | Nagano Pref. | 6600 | Bx & Bm |
| 105 | 7/28 | Nagano Pref. | 70900 | Bx & Bm |
| 106 | 7/28 | Nagano Pref. | 4100 | Bx |
| 107 | 7/29 | Nagano Pref. | 7 | × |
| 108 | 7/30 | Nagano Pref. | 12200 | Bx |
| 109 | 7/30 | Nagano Pref. | 4300 | Bx |
| 110 | 7/30 | Nagano Pref. | 3700 | Bx & Bm |
| 111 | 7/30 | Nagano Pref. | 0 | |
| 112 | 7/30 | Nagano Pref. | 2 | × |
| 113 | 7/30 | Nagano Pref. | 30000 | Bx |
| 114 | 7/30 | Nagano Pref. | 1100 | Bx |
| 115 | 7/30 | Nagano Pref. | 900 | Bx |
| 116 | 7/31 | Nagano Pref. | 13 | Bx |
| 117 | 7/31 | Nagano Pref. | 10000 | Bx & Bm |
| 118 | 8/1 | Nagano Pref. | 4100 | Bx & Bm |
| 119 | 8/4 | Nagano Pref. | 300 | Bx |
| 120 | 8/4 | Nagano Pref. | 5700 | Bx & Bm |
| 121 | 8/5 | Nagano Pref. | 26100 | Bx & Bm |
| 122 | 8/5 | Nagano Pref. | 5200 | Bx & Bm |
| 123 | 8/6 | Nagano Pref. | 4900 | Bx |
| 124 | 8/6 | Nagano Pref. | 500 | Bx |
| 125 | 8/6 | Nagano Pref. | 2300 | Bx |
| 126 | 8/6 | Nagano Pref. | 6800 | Bx |

**Table S7.** Sequences of multiplexed inter-simple sequence repeat genotyping by sequencing (MIG-seq) primer set 1 used for the first polymerase chain reaction (PCR). Names and nucleotide sequences of the forward and reverse primers used in the first PCR of MIG-seq are shown in the 5′–3′ direction.

| Name | Sequence (5'-3') |
|---|---|
| **Forward primers** | |
| $(ACT)_4$TG-f | CGCTCTTCCGATCTCTGACTACTACTACTTG |
| $(CTA)_4$TG-f | CGCTCTTCCGATCTCTGCTACTACTACTATG |
| $(TTG)_4$AC-f | CGCTCTTCCGATCTCTGTTGTTGTTGTTGAC |
| $(GTT)_4$CC-f | CGCTCTTCCGATCTCTGGTTGTTGTTGTTCC |
| $(GTT)_4$TC-f | CGCTCTTCCGATCTCTGGTTGTTGTTGTTTC |
| $(GTG)_4$AC-f | CGCTCTTCCGATCTCTGGTGGTGGTGGTGAC |
| $(GT)_6$TC-f | CGCTCTTCCGATCTCTGGTGTGTGTGTGTTC |
| $(TG)_6$AC-f | CGCTCTTCCGATCTCTGTGTGTGTGTGTGAC |
| **Reverse primers** | |
| $(ACT)_4$TG-r | TGCTCTTCCGATCTGACACTACTACTACTTG |
| $(CTA)_4$TG-r | TGCTCTTCCGATCTGACCTACTACTACTATG |
| $(TTG)_4$AC-r | TGCTCTTCCGATCTGACTTGTTGTTGTTGAC |
| $(GTT)_4$CC-r | TGCTCTTCCGATCTGACGTTGTTGTTGTTCC |
| $(GTT)_4$TC-r | TGCTCTTCCGATCTGACGTTGTTGTTGTTTC |
| $(GTG)_4$AC-r | TGCTCTTCCGATCTGACGTGGTGGTGGTGAC |
| $(GT)_6$TC-r | TGCTCTTCCGATCTGACGTGTGTGTGTGTTC |
| $(TG)_6$AC-r | TGCTCTTCCGATCTGACTGTGTGTGTGTGAC |

**Table S8.** Comparison of classification accuracy levels between theoretical and empirical HIest classification frameworks. Classification outcomes for laboratory individuals of known pedigree that were independent of the parental reference individuals used for HIest estimation are shown for the *B. xylophilus–B. m. kolymensis* and *B. xylophilus–B. m. mucronatus* datasets. Overall and class-specific numbers and percentages of correctly classified individuals were compared between frameworks based on theoretical hybrid index (S)–interspecific heterozygosity (H) positions, empirically estimated class centers and class-specific variation. For the empirical framework, classification accuracy was evaluated using leave-one-out cross-validation, in which each validation individual was classified using empirical class centers and standard deviations calculated after excluding that individual.

| Dataset | Known class | n | Theoretical accuracy (%) | Empirical LOOCV accuracy (%) |
|---|---|---|---|---|
| Bx–Bmk | B1(Bmk) | 23 | 60.9 | 65.2 |
| | B1(Bx) | 5 | 100 | 80 |
| | B2(Bmk) | 24 | 33.3 | 66.7 |
| | F1 | 24 | 87.5 | 100 |
| Bx–Bmk overall | | 76 | 63.2 | 77.6 |
| Bx–Bmm | B1(Bmm) | 4 | 25 | 0 |
| | B2(Bmm) | 16 | 37.5 | 50 |
| | Bmm | 8 | 100 | 25 |
| | Bx | 35 | 100 | 100 |
| | F1 | 32 | 31.3 | 100 |
| Bx–Bmm overall | | 95 | 63.2 | 81.1 |

**Table S9.** HIest-based classification and interpretation of field-collected individuals showing evidence of hybrid ancestry in the *B. xylophilus–B. m. mucronatus* dataset. Primary empirical assignment was based on class-specific standardized distances to empirical reference class centers, with the second-closest class and distance margin shown to indicate classification separation. Ordinary Euclidean (OE) assignment using the same empirical class centers without class-specific standardization is provided as a sensitivity check. Where available, supervised ADMIXTURE ancestry coefficients for *B. xylophilus* [q(Bx)] are also shown. Final interpretation categories conservatively summarize the level of hybrid ancestry supported by these results rather than treating B1/B2 assignments as exact pedigree designations.

| Sample | Year | n SNP (HIest) | S | H | Empirical assignment | Second class | Distance margin | OE sensitivity check | q(Bx) | Interpretation category | Interpretation |
|---|---|---|---|---|---|---|---|---|---|---|---|
| 14-h | 2023 | 29 | 0.58 | 0.66 | F1 | B2(Bmm) | 3.296 | F1 | 0.56 | Strong F1-like | Strong F1-like signal; both methods agree and ADMIXTURE is intermediate. |
| 4-g | 2023 | 26 | 0.56 | 0.58 | F1 | B2(Bmm) | 3.153 | F1 | 0.54 | Strong F1-like | Strong F1-like signal; both methods agree and ADMIXTURE is intermediate. |
| 5-r | 2024 | 32 | 0.53 | 0.67 | F1 | B2(Bmm) | 4.451 | F1 | 0.53 | Strong F1-like | Strong F1-like signal; both methods agree and ADMIXTURE is intermediate. |
| 5-t | 2024 | 39 | 0.54 | 0.62 | F1 | B2(Bmm) | 3.899 | F1 | 0.53 | Strong F1-like | Strong F1-like signal; both methods agree and ADMIXTURE is intermediate. |
| 1-h | 2023 | 24 | 0.59 | 0.24 | B2(Bmm) | B1(Bmm) | 0.954 | F1 | NA | Hybrid, generation uncertain | Hybrid signal supported by both methods; exact generation uncertain (F1 vs backcross-like). |
| 16-h | 2023 | 13 | 0.46 | 0.32 | B2(Bmm) | B1(Bmm) | 0.533 | F1 | NA | Hybrid, generation uncertain | Hybrid-like under both methods, but generation is uncertain and only 13 HIest SNPs were available. |
| 4-i | 2023 | 34 | 0.29 | 0.46 | B1(Bmm) | B2(Bmm) | 0.196 | F1 | 0.22 | Hybrid, generation uncertain | Hybrid signal supported by both methods; F1 vs backcross generation is ambiguous, especially under ordinary distance. |
| 16-e | 2023 | 11 | 0.50 | 0.10 | B2(Bmm) | B1(Bmm) | 0.798 | B1(Bmm) | NA | Backcross-like | Backcross-like signal, but exact class is highly uncertain and only 11 HIest SNPs were available. |
| 3-f | 2023 | 34 | 0.07 | 0.13 | B2(Bmm) | B1(Bmm) | 0.358 | B2(Bmm) | 0 | Backcross-like | Backcross-like toward Bmm; both methods agree on a hybrid class, but B1/B2 resolution is limited. |
| 4-b | 2023 | 23 | 0.08 | 0.15 | B2(Bmm) | B1(Bmm) | 0.319 | B2(Bmm) | NA | Backcross-like | Backcross-like toward Bmm; both methods agree on a hybrid class, but B1/B2 resolution is limited. |
| 12-k | 2024 | 37 | 0.11 | 0.22 | B1(Bmm) | B2(Bmm) | 0.149 | B1(Bmm) | 0.07 | Backcross-like | Backcross-like toward Bmm; both methods agree, but B1 vs B2 is only weakly separated. |
| 15-f | 2024 | 38 | 0.08 | 0.16 | B2(Bmm) | B1(Bmm) | 0.287 | B2(Bmm) | 0.04 | Backcross-like | Backcross-like toward Bmm; both methods agree on a hybrid class, but B1/B2 resolution is limited. |
| 15-i | 2024 | 37 | 0.11 | 0.22 | B1(Bmm) | B2(Bmm) | 0.149 | B1(Bmm) | 0.06 | Backcross-like | Backcross-like toward Bmm; both methods agree, but B1 vs B2 is only weakly separated. |
| 3-f | 2024 | 39 | 0.07 | 0.14 | B2(Bmm) | B1(Bmm) | 0.341 | B2(Bmm) | 0.02 | Backcross-like | Backcross-like toward Bmm; both methods agree on a hybrid class, but B1/B2 resolution is limited. |
| 16-f | 2023 | 12 | 0.68 | 0.17 | B2(Bmm) | B1(Bmm) | 1.199 | Bx | NA | Bx/backcross ambiguous | Ambiguous between Bx-like and backcross-like; method-dependent and based on only 12 HIest SNPs. |
| 16-g | 2023 | 12 | 0.67 | 0.19 | B2(Bmm) | B1(Bmm) | 1.166 | Bx | NA | Bx/backcross ambiguous | Ambiguous between Bx-like and backcross-like; method-dependent and based on only 12 HIest SNPs. |
| 16-i | 2023 | 15 | 0.86 | 0.14 | B2(Bmm) | B1(Bmm) | 1.652 | Bx | NA | Bx/backcross ambiguous | Ambiguous between Bx-like and late backcross-like; method-dependent and based on 15 HIest SNPs. |
| 5-e | 2024 | 43 | 0.92 | 0.16 | B2(Bmm) | Bx | 1.490 | Bx | 0.95 | Bx/backcross ambiguous | Predominantly Bx-like with possible introgression; hybrid assignment is method-dependent and ADMIXTURE strongly favors Bx. |
| 1-i | 2023 | 27 | 0.96 | 0.00 | Bx | B2(Bmm) | 4.789 | Bx | NA | Parental Bx-like | Parental Bx-like by both methods; little HIest support for hybrid ancestry. |
